# A patatin-like phospholipase is crucial for gametocyte induction in the malaria parasite *Plasmodium falciparum*

**DOI:** 10.1101/699363

**Authors:** Ansgar Flammersfeld, Atscharah Panyot, Yoshiki Yamaryo, Philipp Auraß, Jude M. Pryborski, Antje Flieger, Cyrille Botté, Gabriele Pradel

## Abstract

Patatin-like phospholipases (PNPLAs) are highly conserved enzymes of prokaryotic and eukaryotic organisms with major roles in lipid homeostasis. The genome of the malaria parasite *Plasmodium falciparum* encodes four putative PNPLAs with predicted functions during phospholipid degradation. We here investigated the role of one of the plasmodial PNPLAs, a putative PLA_2_ termed PNPLA1, during blood stage replication and gametocyte development. PNPLA1 is present in the asexual and sexual blood stages and here localizes to the cytoplasm. PNPLA1-deficiency due to gene disruption or conditional gene-knockdown had no effect on erythrocytic replication, gametocyte maturation and gametogenesis. However, blood stage parasites lacking PNPLA1 were severely impaired in gametocyte induction, while PNPLA1 overexpression promotes gametocyte formation. The loss of PNPLA1 further leads to transcriptional down-regulation of genes related to gametocytogenesis, including the gene encoding the sexual commitment regulator AP2-G. Additionally, lipidomics of PNPLA1-deficient asexual blood stage parasites revealed overall increased levels of major phospholipids, including phosphatidylcholine (PC), which is a substrate of PLA_2_. Because PC synthesis is pivotal for erythrocytic replication, while the reduced availability of PC precursors drives the parasite into gametocytogenesis, we hypothesize that the high PC levels due to PNPLA1-deficiency prevent the blood stage parasites from entering the sexual pathway.

## 1 INTRODUCTION

Following a first round of multiplication in the human liver, the unicellular malaria parasite *Plasmodium falciparum* enters the blood stream and replicates in human red blood cells (RBCs). The repetitive 48-hours infection cycles, which result in the destruction of the RBC during the release of the newly formed parasites, are responsible for the typical symptoms of malaria such as fever, anemia and organ failure (reviewed in Cowman et al., 2016; Haldar and Mohandas, 2009). Infections with *P. falciparum* are the main cause of the ∼435,000 deaths due to malaria per year, 90% of which occur in the African region (WHO World Malaria Report 2018).

During its intraerythrocytic development, *Plasmodium* induces substantial changes in the structural and functional properties of the RBC. Intraerythrocytic growth leads to an intense period of membrane biogenesis, which includes the formation of external vacuolar transport systems and cell-internal membranous compartments. Membrane synthesis is further crucial for the development of the roughly 16 daughter cells that are formed during erythrocytic schizogony, and these membrane dynamics require synthesis, modification and degradation of phospholipids. While the parasite is capable to *de novo* synthesize the majority of phospholipids, their precursors like the polar heads choline and ethanolamine as well as the fatty acids need to be scavenged from the host blood serum (reviewed in e.g. Ben Mamoun *et al*., 2010; Déchamps, Maynadier, *et al*., 2010; Flammersfeld *et al*., 2018; Vial *et al*., 2003). The total amount of phospholipids increases approximately 5-fold in the infected RBCs with phosphatidylcholine (PC) and phosphatidylethanolamine (PE) being the main membrane components (Beaumelle & Vial, 1988; Botté et al., 2013; Gulati et al., 2015; Simões, Roelofsen, & Op den Kamp, 1992; reviewed in Déchamps, Maynadier, et al., 2010).

During the erythrocytic infection cycle, a proportion of blood stage parasites enter the sexual pathway, which results in the production of the transmissible gametocytes. Gametocyte induction is triggered in response to stress factors and particularly depends on the availability of PC precursors in the human serum, i.e. low levels of serum-derived lysoPC and choline cause the blood stage parasites to enter the sexual pathway (Brancucci et al., 2017; Wein et al., 2018). These gametocytes, once matured, transform into gametes following their uptake by blood-feeding *Anopheles* mosquitoes. Gametogenesis is followed by sexual reproduction of the parasite, an event preluding the mosquito-specific phase of the plasmodial life-cycle (reviewed in Bennink, Kiesow, & Pradel, 2016; Kuehn and Pradel, 2010).

Membrane biogenesis in the intraerythrocytic blood stages requires the action of phospholipases, lipolytic enzymes that are classified into four groups, A, B, C and D corresponding to the hydrolysis activity (reviewed in Déchamps, Shastri, *et al*., 2010; Flammersfeld *et al*., 2018; Vial and Ben Mamoun, 2005). A recent analysis of the *P. falciparum* genome identified 22 putative phospholipases. The main proportion of these enzymes exhibit a predicted α/β hydrolase domain and 12 of these were annotated as lysophospholipases (Flammersfeld et al., 2018). In addition, four putative patatin-like phospholipases (PNPLAs), lipolytic enzymes that display lipase and transacylase properties, are encoded in the *P. falciparum* genome. PNPLAs can typically be found both in eukaryotes and in a variety of bacteria, where they have major roles in lipid homeostasis, but where they can also be involved in membrane degradation and cell signaling (reviewed in Banerji & Flieger, 2004; Kienesberger, Oberer, Lass, & Zechner, 2009). They share conserved common protein domains with patatin, a storage glycoprotein of potato tuber with lipid acyl hydrolase activity (Senda *et al*., 1996; reviewed in Shewry, 2003). All PNPLAs exhibit a catalytic Ser-Asp dyad with the serine residue being embedded within the conserved penta-peptide Gly-Xaa-Ser-Xaa-Gly (reviewed in Arpigny and Jaeger, 1999; Banerji and Flieger, 2004; Ramanadham *et al*., 2015). Via their phospholipase A_2_ (PLA_2_) activity, PNPLAs are able to generate lysophospholipids and fatty acids, including polyunsaturated fatty acids, which are either further processed within the lipid metabolism pathway or which may act as signaling molecules (Lévêque et al., 2017; reviewed in Kienesberger *et al*., 2009; Ramanadham *et al*., 2015; Wilson and Knoll, 2018).

We here report on the functional characterization of one of the four plasmodial PNPLAs, termed PNPLA1. The protein is found in the cytosol of the asexual and sexual blood stages of *P. falciparum*. Asexual blood stages lacking PNPLA1 exhibit upregulated phospholipid levels as well as impaired gametocyte induction both under normal growth conditions and following growth in phospholipid precursor-deficient medium, suggesting that the deregulated phospholipid levels in the asexual blood stages render these less sensitive to triggers of gametocytogenesis.

## 2 RESULTS

### 2.1 PNPLA1 is a patatin-like phospholipase of the *P. falciparum* blood stages

The gene PF3D7_0209100 of *P. falciparum* encodes a putative patatin-like phospholipase, henceforth termed PNPLA1, with a molecular weight of 78 kDa. PNPLA1 is predicted to be a PLA_2_ with a patatin-like phospholipase domain spanning from aa 338 to aa 544 and a C-terminal FabD/lysophospholipase-like domain ranging from aa 332 to aa 673 (Fig. 1A). The characteristic Gly-Xaa-Ser-Xaa-Gly motif is located between aa 381 and aa 385, while the catalytic aspartate of the dyad is located at aa 531. 3D modelling of PNPLA1 was done using the I-TASSER server based on the crystal structure of PDB hit 6AUN (Malley et al., 2018), a human calcium-independent phospholipase A_2_-β with cytosolic localization that is involved in diverse biological processes including fat catabolism, cell differentiation and phospholipid remodeling (reviewed in Barbour and Ramanadham, 2017) (Fig. S1A). The model shows the serine (aa 383) and aspartate (aa 531), representing the catalytic dyad, being in close spatial proximity within the C-terminal catalytic domain. Phylogenetic analysis of PNPLA1 and related proteins from other plasmodia identified the highest identities with the homologous proteins of *P. ovale* and *P. malariae*, while the highest diversity was found in the PNPLA1 homolog of *P. berghei* (Fig. S1B). All of the plasmodial PNPLA1 homologs share the GXSXG motif and the catalytic aspartate; the homologs of *P. vivax* and *P. knowlesi* further exhibit N-terminal extensions (Fig. S2).

**FIGURE 1.**
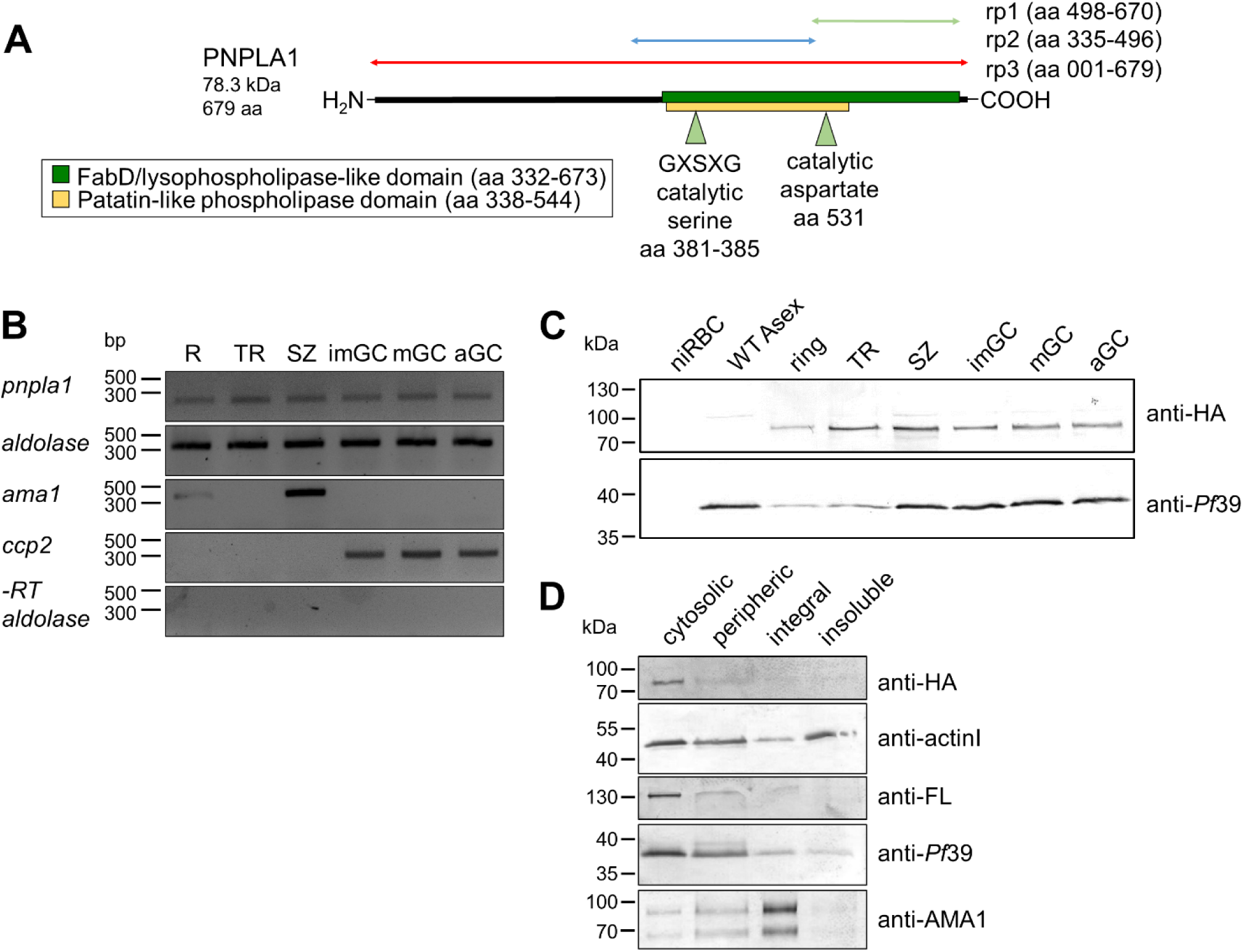
PNPLA1 is expressed in the asexual blood and gametocyte stages of *P. falciparum*. **(A)** Schematic depicting the domain architecture of PNPLA1. The FabD/lysophospholipase-like domain (green) and the patatin-like phospholipase domain (yellow), the catalytic serine lipase motif Gly-Xaa-Ser-Xaa and the catalytic aspartate, both constituting the serine-aspartate dyad, are highlighted. Regions used for generation of recombinant proteins (rp) are indicated. aa, amino acids; FabD, fatty acid binding domain; G, glycine; S, serine. **(B)** Analysis of *pnpla1* transcription. Diagnostic RT-PCR was used to amplify *pnpla1* transcript (220 bp) from cDNA generated from total RNA of rings (R), trophozoites (TR), schizonts (SZ), as well as immature (GCII-IV), mature (GCV) and activated gametocytes (GC 15’ p.a.). Transcript analysis of *ama1* (407 bp) and *ccp2* (286 bp) were used to demonstrate purity of the asexual and sexual blood stage samples. Transcript analysis of *aldolase* (378 bp) was used as loading control, RNA samples lacking reverse transcriptase (-RT) were used to prove absence of gDNA. **(C)** PNPLA1-HA expression in blood stage parasites. Western blotting of lysates from rings, trophozoites (TR), schizonts (SZ), immature (imGC), mature (mGC) and activated gametocytes (aGC, 30 min p.a.) of *Pf*PNPLA1-HA, using rabbit anti-HA antibody, detected PNPLA1-HA at the expected size of ∼78 kDa. Lysates of mixed WT asexual blood stages (WT Asex) and non-infected RBCs (niRBCs) were used as negative controls; equal loading was confirmed using polyclonal mouse antisera against *Pf*39 (∼39 kDa). **(D)** Subcellular localization of PNPLA1-HA. Lysates of the PNPLA1-HA line were used to extract soluble, integral, peripheric and unsoluble protein fractions. Samples were immunoblotted with rabbit anti-HA antibodies to detect PNPLA1-HA (∼82 kDa); immunoblotting with mouse antisera against actin I (∼42 kDa), falcilysin (∼138 kDa), *Pf*39 (∼39 kDa), and AMA1 (∼66 and 88 kDa) were used for fraction controls. Results are representative of two to three independent experiments.

Diagnostic RT-PCR was performed to investigate expression of *pnpla1* in the *P. falciparum* blood stages and demonstrated transcripts in the ring and trophozoite stages and in schizonts. Similar transcript levels were detected in immature (stage II-IV) and mature (stage V) gametocytes as well as in gametocytes at 15’ post-activation (p.a.) (Fig 1B). Transcript analysis of the housekeeping gene *aldolase* was used as a loading control, and purity of the asexual blood stage and gametocyte samples was demonstrated by amplification of transcripts for the asexual blood stage-specific gene *pfama1* (apical membrane antigen 1) (Peterson et al., 1989) and for the gametocyte-specific gene *pfccp2* (LCCL-domain protein 2) (Pradel et al., 2004). The cDNA preparations lacking the reverse transcriptase (-RT) were used to verify that samples were devoid of gDNA (Fig. 1B).

Mouse antisera were generated against two recombinant PNPLA1 peptides comprising parts of the lysophospholipase and the patatin-like phospholipase domain (PNPLA1rp1 and PNPLA1rp2, respectively; see Fig 1A) to be used for protein expression analysis. In addition, a full-length recombinant protein was used in WB to confirm the specificity of PNPLA1rp1 and PNPLA1rp2 (Fig. S3A). Indirect immunofluorescence assays (IFAs), using either of the two antisera, revealed that PNPLA1 was abundantly expressed in the asexual blood stages, where it localized to the cytosol (Figs 2A; S3B). PNPLA1 was further detected in the cytosol of gametocytes during development and at 15 min and 30 min p.a. (Fig. 2B; S3C). Counterlabelling with antibodies either directed against Pfs25, a female-specific EGF domain protein, or against Pfs230, a cysteine-rich protein present in both male and female gametocytes (reviewed in Pradel, 2007), demonstrated that PNPLA1 is expressed in a gender-independent pattern (Fig S3D). When sera from non-immunized mice (NMS) was used in the IFAs as a negative control, no labelling was detectable in either asexual blood stages or gametocytes (Fig. S3E).

**FIGURE 2.**
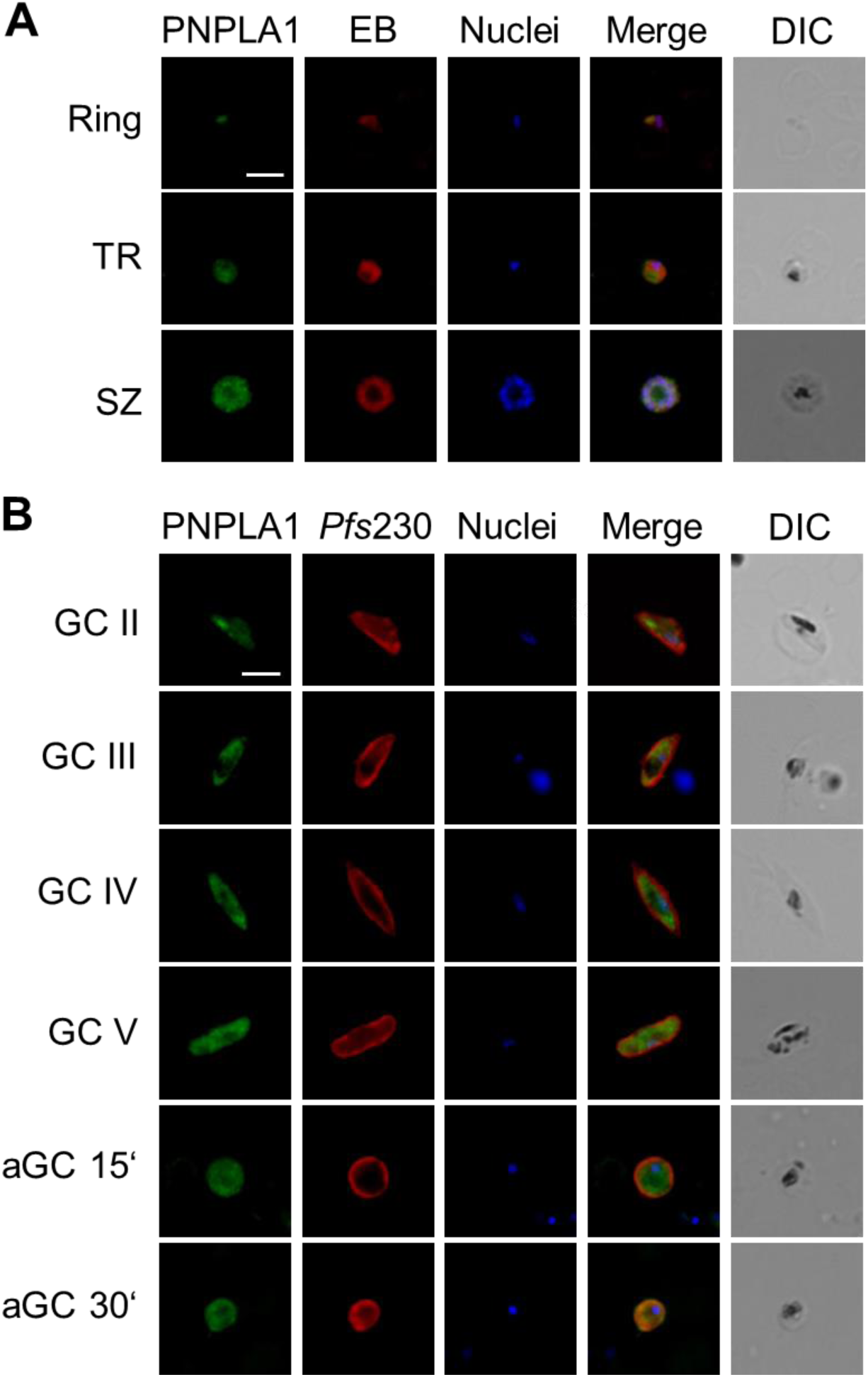
PNPLA1 localizes to the blood stage cytosol. Rings, trophozoites (TR), schizonts (SZ), gametocytes (GCII-GCV) and activated gametocytes (aGC; at 15 and 30 min p.a.) of WT were immunolabelled with mouse anti-PNPLA1rp1 antisera (green). The asexual blood stages were stained with EB; the gametocytes were counterlabelled with rabbit anti-*Pfs*230 sera (red). The parasite nuclei were highlighted by Hoechst33342 nuclear stain (blue). DIC, differential interference contrast. Bar, 5 µm. Results are representative of three independent experiments.

### 2.2 PNPLA1-deficient blood stage parasites are impaired in gametocyte induction

We generated two PNPLA1-deficient lines for functional analyses. Firstly, *P. falciparum* parasites lacking the *pnpla1*-encoding gene were generated via single cross-over homologous recombination, using the vector pCAM-BSD (Fig. S4A). Two clonal lines (A12 and C11) of the PNPLA1-knock out (KO) were isolated and successful integration of the plasmid into the genome was verified by diagnostic PCR and by sequencing of the integration loci (Fig. S4B, S5). The absence of the protein in PNPLA1-KO schizonts of both lines compared to the wildtype (WT) was subsequently demonstrated by IFA (Fig. S4C). Similarly, WB using the mouse anti-PNPLA1rp1 antisera, could detect a PNPLA1-positive protein band in blood stage lysates of the WT, but not in lysates of the two PNPLA1-KO lines (Fig. S4D).

In addition, we generated a conditional PNPLA1-knock down (KD) line (using the pARL-HA-*glmS* vector), by fusing the sequences coding for a hemagglutinin A tag and for the *glmS* element to the 3’-region of *pnpla1* (Fig. S6A). In the PNPLA1-KD, the *pnpla1* mRNA is expressed under the control of the *glmS* ribozyme (Prommana et al., 2013), which catalyzes its own cleavage in the presence of glucosamine (GlcN) and thus triggers degradation of the *pnpla1* mRNA. Vector integration was confirmed by diagnostic PCR and sequencing of the integration sites (Fig. S6A, B; S7). Immunolabelling with an anti-HA antibody confirmed the synthesis of a HA-tagged PNPLA1 fusion protein in the untreated PNPLA1-KD line, while no signal was detected in WT (Fig. S4C). Similarly, immunoblotting with the anti-HA antibody detected the PNPLA1 fusion protein running with a molecular weight of 82.2 kDa and confirmed protein expression in the asexual blood stages as well in gametocytes during development and following activation (Fig. 1C). Subcellular fractioning, using an anti-HA antibody, further demonstrated that PNPLA1-HA locates to the cytosolic fraction of the blood stage parasite, while the protein was absent in the peripheric, the integral and the insoluble factions (Fig. 1D). Immunoblotting with antibodies against the cytoskeletal element actin1 (Ngwa *et al*., 2013; reviewed in Baum *et al*., 2008;) the cytosolic protease falcilysin (Weißbach, Golzmann, Bennink, Pradel, & Ngwa, 2017), the endoplasmic reticulum-associated protein *Pf*39 (Simon et al., 2009) and the transmembrane protein AMA1 (Boes et al., 2015; Peterson et al., 1989) were used for fraction controls. To verify the conditional down-regulation of PNPLA1 synthesis, asexual blood stage parasites were treated with 2.5 mM GlcN for 72 h. Upon GlcN addition, PNPLA1 levels were significantly reduced to 40.00 ± 13.4% compared to the untreated control (Fig. S6D, E).

Initial phenotype analyses of the two PNPLA1-KO lines showed that intraerythrocytic development of the asexual blood stages was normal compared to WT and no differences in parasitemia over a period of 120 h as well as in blood stage morphology and progression and in the numbers of merozoites formed per schizont were observed (Fig. 3A-C; S8A). Furthermore, no significant differences in the exflagellation behavior as well as in the formation of macrogametes and zygotes following in vitro activation of the PNPLA1-KO gametocytes was seen (Fig. 3D). When gametocytemia was compared between the PNPLA1-KO lines and the WT over a period of 14 d, however, significant reductions in the numbers of gametocytes were documented (Fig. 3E).

**FIGURE 3.**
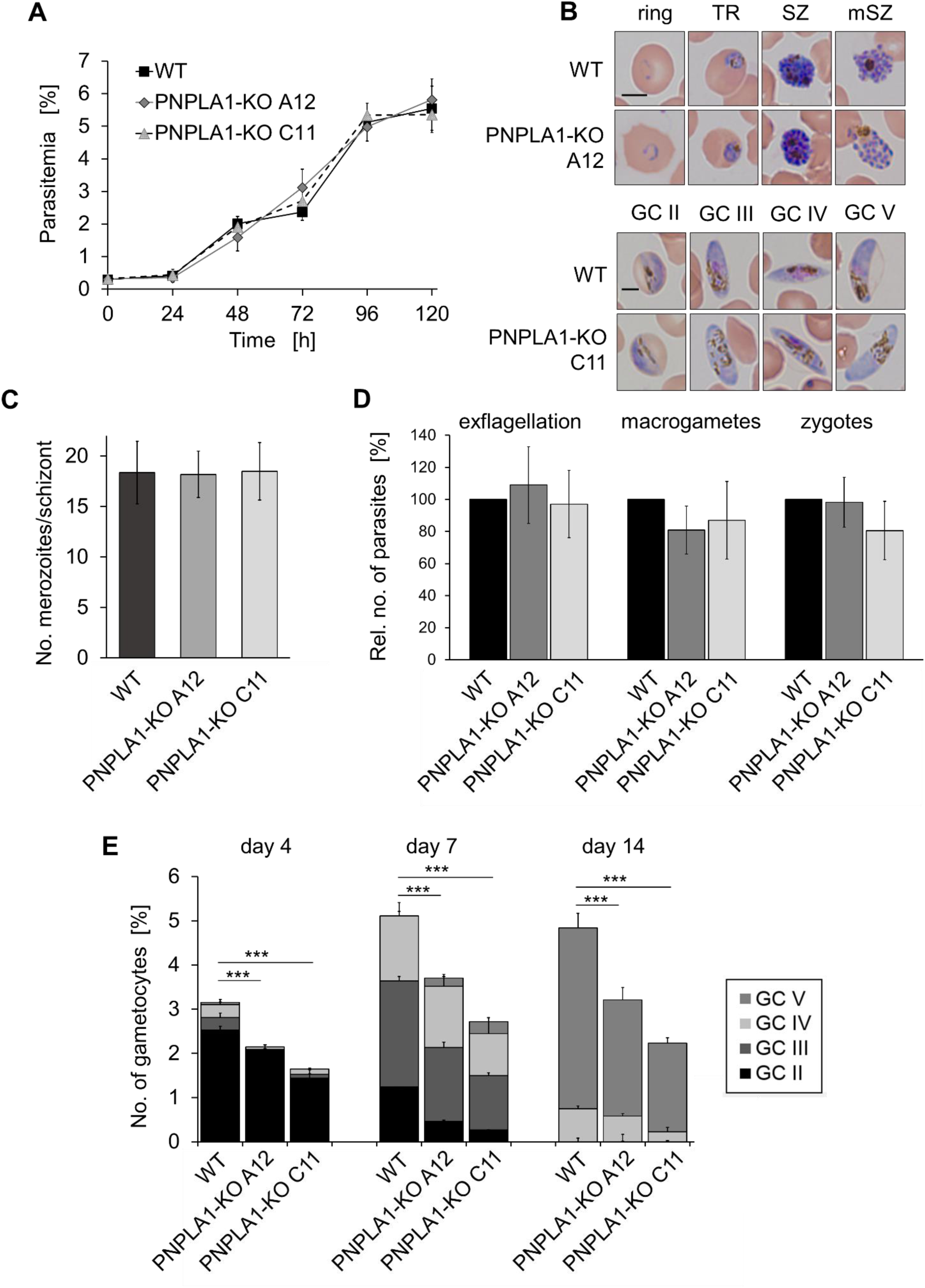
PNPLA1-KO parasites are impaired in gametocytogenesis. **(A)** Asexual blood stage replication of the PNPLA1-KO. Ring stage cultures of WT and PNPLA1-KO A12 and C11 with a starting parasitemia of 0.25% were maintained in cell culture medium and parasitemia was followed via Giemsa smears over a time-period of 0-120 h. The experiment was performed in triplicate (mean ± SD). **(B)** Morphology of the PNPLA1-KO blood stages. The morphology was demonstrated via Giemsa staining of asexual blood stages and gametocytes of the PNPLA1-KO A12 and C11 lines and WT. TR, trophozoites; SZ, schizonts; mSZ, mature schizonts; GC, gametocytes II-V. Bar, 5 µm. **(C)** Merozoite formation in PNPLA1-KO schizonts. Schizonts were stained with EB and merozoites were highlighted by Hoechst33342 nuclear stain. Merozoite numbers were counted in triplicate for 75 schizonts per parasite line. Data are shown as mean ± SD. **(D)** Gametogenesis of the PNPLA1-KO. Mature WT and PNPLA1-KO A12 and C11 gametocytes were activated *in vitro*. Samples were taken at 15 min p.a. (exflagellation centers), 30 min p.a. (macrogametes) and 6 h p.a. (zygotes). Exflagellation centers were counted in 30 optical fields for four times using light microscopy. Zygotes and macrogametes were immunolabelled with anti-*Pfs*25 antibody and counted in 30 optical fields in triplicate. Results are shown as mean ± SD (WT set to 100%). **(E)** Gametocyte development of the PNPLA1-KO. WT and the PNPLA1-KO lines A12 and C11 with a starting parasitemia of 2.0% were grown in cell culture medium and numbers of gametocytes were followed via Giemsa smears over a time-period of 14 d. The experiment was performed in triplicate (mean ± SD). ***p≤0.001 (One-Way ANOVA with Post-Hoc Bonferroni Multiple Comparison test). The data (A-E) are representative of three to five independent experiments.

To investigate the effect of PNPLA1-deficiency on gametocytes in more detail, we used the PNPLA1-KD line in the follow-up experiments. We confirmed that reduced PNPLA1 synthesis did not lead to any differences in parasitemia over a period of 120 h as well as in blood stage morphology and progression, when these were treated or not treated with GlcN (Fig. 4A; S8B, S9). The low PNPLA1-levels, however, significantly reduced gametocyte numbers in the GlcN-treated PNPLA1-KD line, when this line was compared to untreated PNPLA1-KD parasites and treated or untreated WT parasites (Fig. 4B). To determine, if the PNPLA1-deficiency effects gametocyte induction or rather early gametocyte development, we treated the blood stages with GlcN either before or after inducing sexual commitment by the addition of lysed RBCs. We observed that PNPLA1 down-regulation prior to gametocyte induction resulted in significantly reduced gametocyte numbers, while PNPLA1 down-regulation after gametocyte induction had no effect (Fig. 4C). Untreated PNPLA1-KD parasites as well as GlcN-treated and untreated WT were used for controls in these experiments.

**FIGURE 4.**
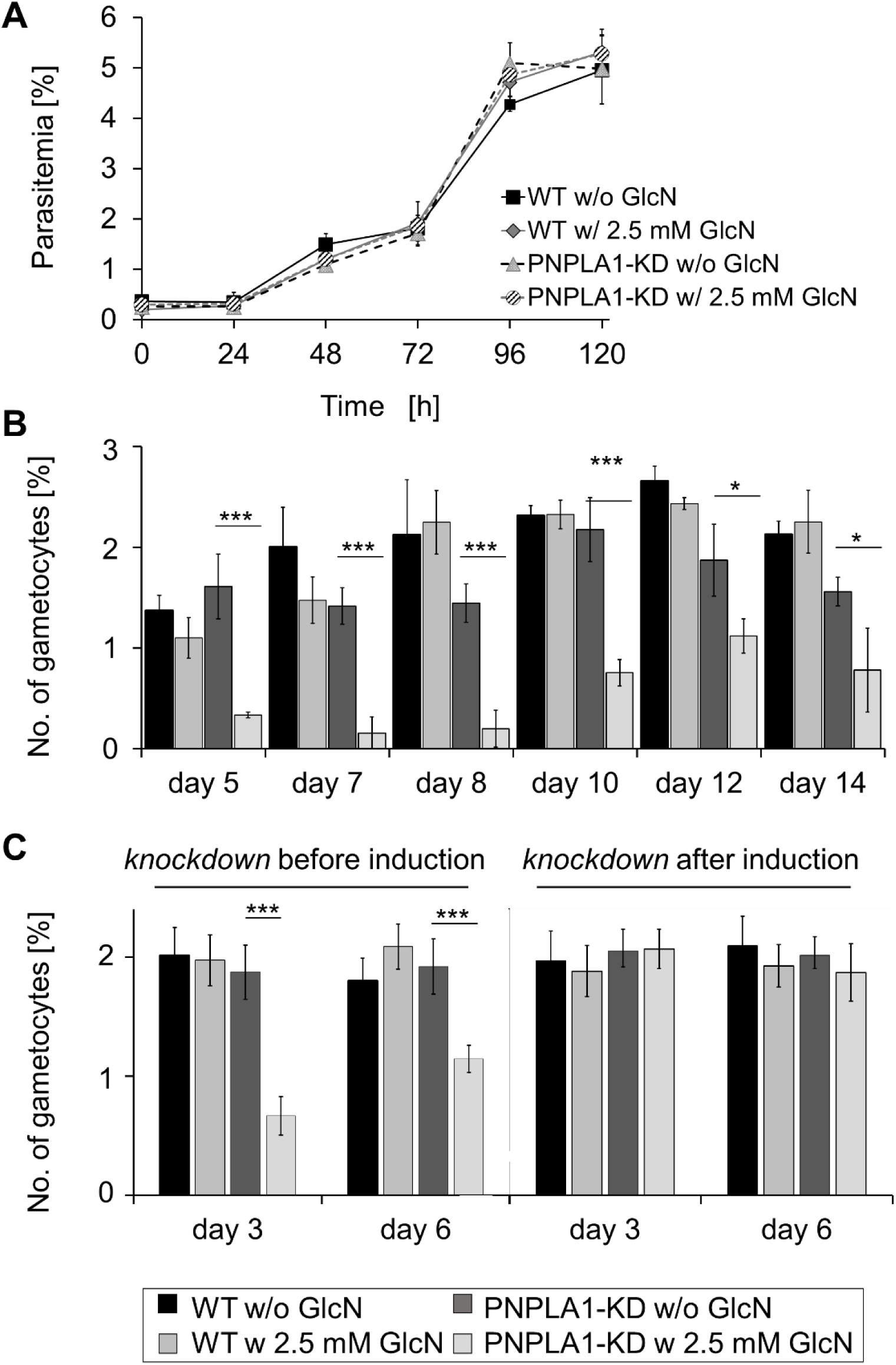
PNPLA1-KD parasites are impaired in gametocyte induction. **(A)** Asexual blood stage replication of the PNPLA1-KD. Ring stage cultures of WT and the PNPLA1-KD with a starting parasitemia of 0.25% were maintained in cell culture medium supplemented with 2.5 mM GlcN to knockdown PNPLA1 protein expression and parasitemia was followed via Giemsa smears over a time-period of 0-120 h. Untreated (without GlcN) cultures and WT were used as controls. The experiment was performed in triplicate (mean ± SD). **(B)** Gametocyte formation of the PNPLA1-KD. WT and PNPLA1-KD parasites with a starting parasitemia of 2.0% were grown in cell culture medium supplemented with 2.5 mM GlcN and numbers of gametocytes were followed via Giemsa smears over a time-period of 14 d. Untreated (without GlcN) cultures and WT were used as controls. The experiment was performed in triplicate (mean ± SD). ***p≤0.001 (One-Way ANOVA with Post-Hoc Bonferroni Multiple Comparison test). **(C)** Gametocyte induction and maturation of the PNPLA1-KD. WT and PNPLA1-KD parasites with a starting parasitemia of 2.0% were grown in cell culture medium until a parasitemia of 10.0% was reached. For gametocyte induction, cultures were exposed to lysed RBCs for 24 h, washed with cell culture medium, and further grown in cell culture medium supplemented with 50 mM GlcNac to kill the asexual blood stages. Cultures were treated either before or after gametocyte induction with 2.5 mM GlcN to knockdown PNPLA1 protein expression. Numbers of gametocytes were determined via Giemsa smears on day 3 and 6 p.g.i. The experiment was performed in triplicate (mean ± SD). ***p≤0.001 (One-Way ANOVA with Post-Hoc Bonferroni Multiple Comparison test). The data (A-C) are representative of two to three independent experiments.

### 2.3 PNPLA1 deficiency counteracts gametocyte induction by serum-free medium

In a recent study, Brancucci et al. (2017) reported that serum-free cell culture medium (-SerM), which only contained the minimally required fatty acid species C16:0 and C18:1 necessary for the growth of *P. falciparum* blood stages (Mi-Ichi, Kano, & Mitamura, 2007; Mi-Ichi, Kita, & Mitamura, 2006), triggers sexual commitment. This effect could be reverted by addition of lysoPC to the serum. We thus repeated the above experiments and induced gametocyte formation by cultivation of the blood stage parasites in –SerM. We confirmed that in the WT treatment with –SerM resulted in an increased formation of gametocytes compared to medium supplemented with lysoPC, choline or human serum (Fig. 5A). The two PNPLA1-KO lines, however, did not respond to –SerM and gametocyte numbers were significantly reduced (Fig. 5A). Similarly, the PNPLA1-KD line did not respond to –SerM, when treated with GlcN prior to gametocyte induction, while the untreated PNPLA1-KD line formed higher numbers of gametocytes in –SerM compared to medium supplemented with lysoPC, choline or human serum (Fig. 5B).

**FIGURE 5.**
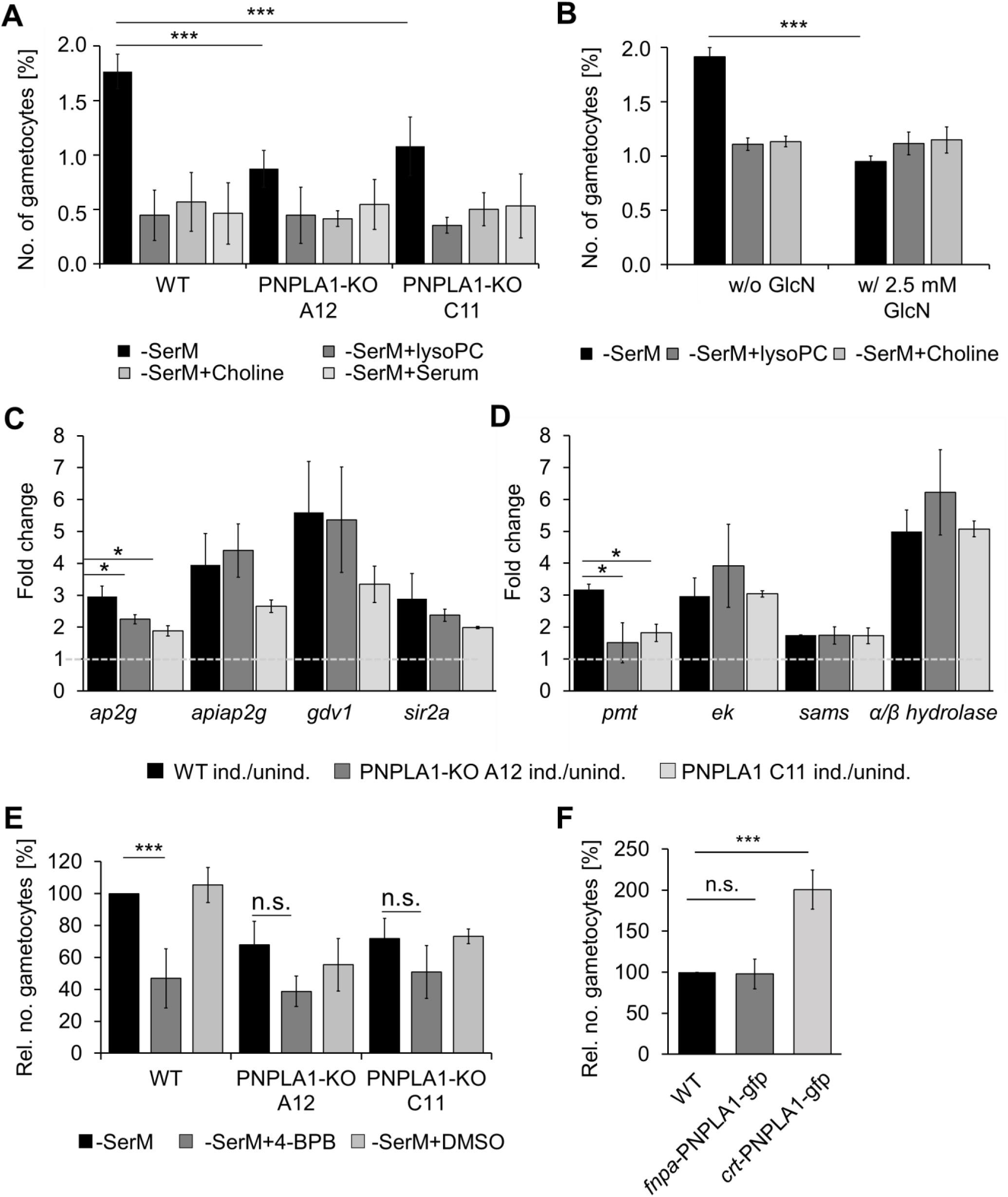
PNPLA1 levels effect gametocyte induction due to serum-depleted medium. **(A, B)** Gametocyte induction in the PNPLA1-deficient lines. Tightly synchronized parasite cultures of WT and the PNPLA1-KO lines A12 and C11 (**A**) or the PNPLA1-KD line (**B**) were treated with -SerM at 28 ± 2 h.p.i for 22 h and further grown in cell culture medium supplemented with 50 mM GlcNac for 5 d to kill the asexual parasites. For control, -SerM was supplemented with 20 µM lysoPC, with 300 µM choline and 11.1 mM glucose, or with 10% v/v serum. Numbers of gametocytes were determined via Giemsa smears on day 5 p.g.i. The experiments (A, B) were performed in triplicate (mean ± SD). **(C, D)** Transcription in the PNPLA1-KO following gametocyte induction. Tightly synchronized parasite cultures of WT and the PNPLA1-KO lines A12 and C11 were treated with -SerM at 28 ± 2 h.p.i. and total RNA was collected at 46 ± 2 h.p.i. Transcript levels for four genes encoding transcriptional regulators of sexual commitment (**C**) and enzymes of the phospholipid metabolic pathway (**D**) were determined by real-time RT-PCR. The Ct values were normalized with the Ct values of the gene encoding the seryl tRNA-ligase (PF3D7_0717700) as reference. The quotients of the fold changes between the induced and non-induced cultures of WT, PNPLA1-KO A12 and PNPLA1-KO C11 are shown. Measurements were performed in triplicate from three independent biological cDNA preparations (mean ± SD). ***p≤0.001 (One-Way ANOVA with Post-Hoc Bonferroni Multiple Comparison test). *ap2g*, transcription factor with ap2 domains; *apiap2g*, transcription factor with ap2 domains; *gdv1*, gametocyte development protein 1; *sir2a*, transcriptional regulatory protein sir2a; *pmt*, phosphoethanolamine N-methyltransferase; *ek*, ethanolamine kinase; *sams*, S-adenosylmethionine synthetase. **(E)** Effect of PLA_2_ inhibitor 4-BPB on gametocyte induction. WT and the PNPLA1-KO lines A12 and C11 treated with -SerM supplemented with 4-BPB at IC_50_ concentration at 28 ± 2 h.p.i for 22 h and further grown in cell culture medium supplemented with 50 mM GlcNac for 5 d to kill the asexual parasites. Parasites cultivated in medium with or without 1% v/v DMSO were used as control. **(F)** Effect of PNPLA1 overexpression on gametocyte development. Tightly synchronized parasite cultures of WT and the two lines *fnpa*-PNPLA1-GFP and *crt*-PNPLA1-GFP were treated with -SerM at 28 ± 2 h.p.i for 22 h and further grown in cell culture medium supplemented with 50 mM GlcNac for 5 d to kill the asexual parasites. Numbers of gametocytes were determined via Giemsa smears on day 5 p.g.i. The experiments (E, F) were performed in triplicate (mean ± SD; WT set to 100%). ***p≤0.001 (One-Way ANOVA with Post-Hoc Bonferroni Multiple Comparison test). The data (A-F) are representative for three to five independent experiments.

Brancucci et al. (2017) further described that lysoPC-depletion triggers an immediate transcriptional upregulation of a variety of genes, including such encoding transcriptional regulators of sexual commitment like *ap2-g* (Brancucci et al., 2017; Kafsack et al., 2014; Sinha et al., 2014) and essential gametocytogenesis factors like *gdv1* (Brancucci et al., 2017; Eksi et al., 2012). Furthermore, genes coding for enzymes that are known to be involved in the utilization of ethanolamine to generate PC were upregulated (Bobenchik et al., 2010; Brancucci et al., 2017; Pessi, Choi, Reynolds, Voelker, & Mamoun, 2005; Pessi, Kociubinski, & Mamoun, 2004). We chose eight genes that were found to be upregulated under lysoPC-limiting conditions and investigated their transcriptional levels at 46 ± 2 h.p.i. (hours post-invasion) to compare induced to non-induced controls. For all of the eight genes, a 2- to 5-fold transcriptional upregulation could be observed in the –SerM-cultivated WT (Fig. 5C, D). A similar transcriptional up-regulation was seen in the PNPLA1-KO lines A12 and C11, but the fold-changes were decreased. For *ap2-g*, and the phosphoethanolamine N-methyltransferase PMT that catalyzes the conversion of PE into PC, significantly reduced transcript levels compared to the –SerM-induced WT control were observed.

In a next step, the effect of commercial PLA_2_ inhibitors on gametocyte induction was tested. First, the antimalarial effect of the four inhibitors 4-bromophenacyl bromide (4-BPB), bromoenol lactone, ASB14780 and N-(p-amylcinnamoyl) anthranilic acid on erythrocytic replication was investigated, using the Malstat assay. The highest antimalarial effect was detected for 4-BPB with an IC_50_ value of 5.6 ± 0.82 µM (Table 1). This concentration was used to treat asexual blood stages when inducing gametocytes by treatment with –SerM. When the asexual blood stages were treated with 4-BPB, significantly reduced gametocyte numbers were observed in the WT upon –SerM incubation compared the untreated WT and the solvent control (Fig. 5E). On the other hand, in the two PNPLA1-KO lines, no significant differences were observed in 4-BPB-treated parasites compared to the controls. 4-BPB treatment, however, had no significant effect on gametocyte maturation and gametocyte stage progression, when added to early stage II gametocytes, while epoxomicin used for positive control (Aminake et al., 2011) killed the gametocytes (Fig. S10A, B).

**TABLE 1.**
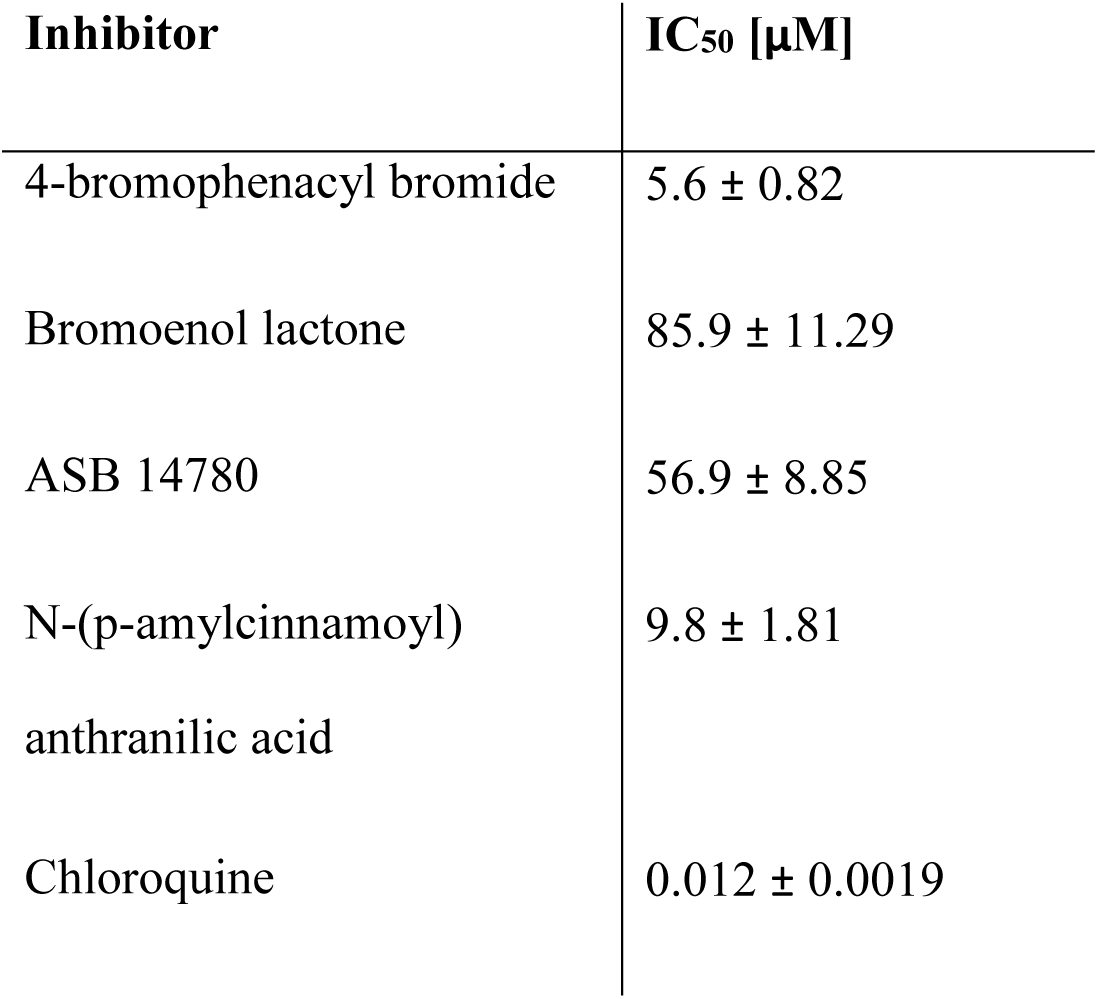
Antimalarial activities of PLA_2_ inhibitors.

We also investigated the reverse effect, i.e. the effect of PNPLA1 overexpression on gametocyte induction. The sequence for full-length PNPLA1 was cloned into the vector pARL-GFP, which was driven either under the control of the ubiquitously active *crt* promotor (Külzer et al., 2010; Przyborski et al., 2005; Spork et al., 2009) or the gametocyte-specific *fnpa* promotor (Bennink et al., 2018; Pradel et al., 2004) (Fig. S11A). Presence of the plasmids in the transfectants was verified by diagnostic PCR, purified plasmid DNA was used for control (Fig. S11B). Live imaging of corresponding lines showed that the *crt* promotor allowed the episomal expression of PNPLA1 in the transfected asexual blood stages (line *crt*-PNPLA1-GFP), while the protein was expressed in transfected gametocytes when controlled by the *fnpa* promotor (line *fnpa*-PNPLA1-gfp) (Fig. S11C). WT parasites did not show any GFP expression (Fig. S11D). Asexual blood stages of the transfectant lines *crt*-PNPLA1-GFP and *fnpa*-PNPLA1-GFP were subsequently induced by –SerM, which resulted in significantly increased gametocyte numbers in the *crt*-PNPLA1-GFP line at day 5 post-gametocyte induction (p.g.i.) compared to the WT and *fnpa*-PNPLA1-GFP (Fig. 5F). No differences in the morphology of the asexual and sexual blood stages, though, were observed in lines *crt*-PNPLA1-GFP and *fnpa*-PNPLA1-GFP compared to WT (Fig. S12).

### 2.4 PNPLA1-deficiency results in an overall increase of major phospholipids

Since the deficiency of PNPLA1 impaired the proper gametocytogenesis pathway, we investigated whether the parasite lipid composition was altered in the PNPLA1-KO lines, accordingly to the putative role of the enzyme and the role of phospholipids for maintaining normal asexual blood stage growth. Total lipids from synchronized ring stages of WT and the PNPLA1-KO lines A12 and C11 were extracted and mixed with an internal lipid standard mix (LIPDOMIX standard, Avanti). Each lipid class was separated by high performance thin layer chromatography and then extracted from the silica gel. Total fatty acids from each lipid class were methanolized to fatty acid methyl ester and then quantified by GC-MS. The general phospholipid levels of the two PNPLA1 lines was slightly higher compared to the WT (Fig. 6A). This increase was not particularly associated to one specific lipid class but more to general lipid increase. The phospholipid changes, however, were particularly drastic for PC, which was significantly increased in the PNPLA1 lines compared to the WT (Fig.6 B). Noteworthy, he difference in PC levels did not alter the overall phospholipid composition (Fig.6C), probably for the parasite to keep its membrane integrity. Solely for lysobisphosphatidic acid (LBPA) levels, a decrease in the PNPLA1-KO lines compared to WT could be observed (Fig. 6B, C). The lysoPC content in the different lines was below detection level.

**FIGURE 6.**
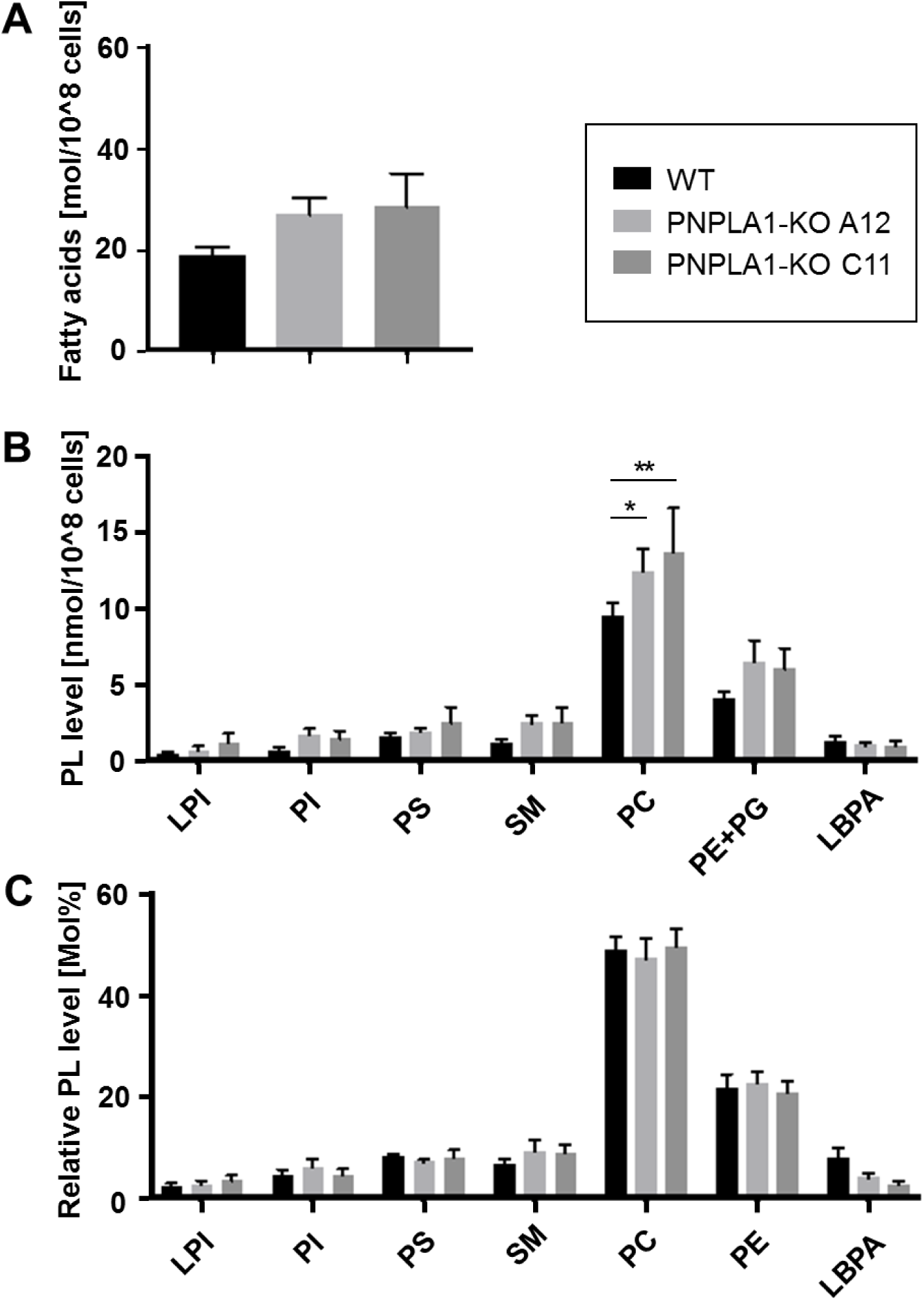
PNPLA1-KO parasites exhibit altered phospholipid levels. **(A)** PNPLA1-KO parasites have increased fatty acid contents. Lipids were extracted from WT and the PNPLA1-KO lines A12 and C11 and employed to lipidomics. Fatty acid contents were compared after the normalization to parasite number of 10^8^. (**B**) PNPLA1-KO parasites exhibit altered levels of phospholipids. The contents of selected phospholipids were compared following normalization. Statistic analysis was performed with GraphPad Prism 5 software using two-way ANOVA analysis. P value is shown as *, p<0.05; **, p<0.01. (**C**) PNPLA1-KO parasites are not altered in the relative phospholipid levels. The relative phospholipid contents are shown as Mol%. PC, phosphatidylcholine; PE, phosphatidylethanolamine; PG, phosphatidylglycerol; PI, phosphatidylinositol; PL, phospholipid; PS, phosphatidylserine; LPI, lysophosphatidylinositol; LBPA, lysobisphosphatidic acid; SM, sphingomyelin. The lipidomic analysis were performed using four biological replicates of each line.

## 3 DISCUSSION

During infection with *P. falciparum*, gametocytes begin to develop approximately one week after the appearance of parasites in the human blood. A small proportion of committed parasites leaves the asexual blood stage cycle and enters the sexual pathway on a continuous basis, an event probably driven by endogenous factors. The gametocyte commitment rate, however, increases drastically in response to environmental stress signals, including parasite density, anaemia, host immune response or drug treatment (reviewed in Alano, 2007; Kuehn and Pradel, 2010). Increasing data suggest that the lack of phospholipid precursors in the human serum triggers gametocyte induction. Particularly, low serum levels of lysoPC, which was recently shown to act as the main exogenous precursor of PC, cause the blood stage parasites to enter the sexual pathway (Brancucci et al., 2017; Wein et al., 2018). Despite previous assumptions that choline is acquired from serum, new data indicate that the majority of choline to be incorporated in PC is generated from imported lysoPC (Fig. 7).

**FIGURE 7.**
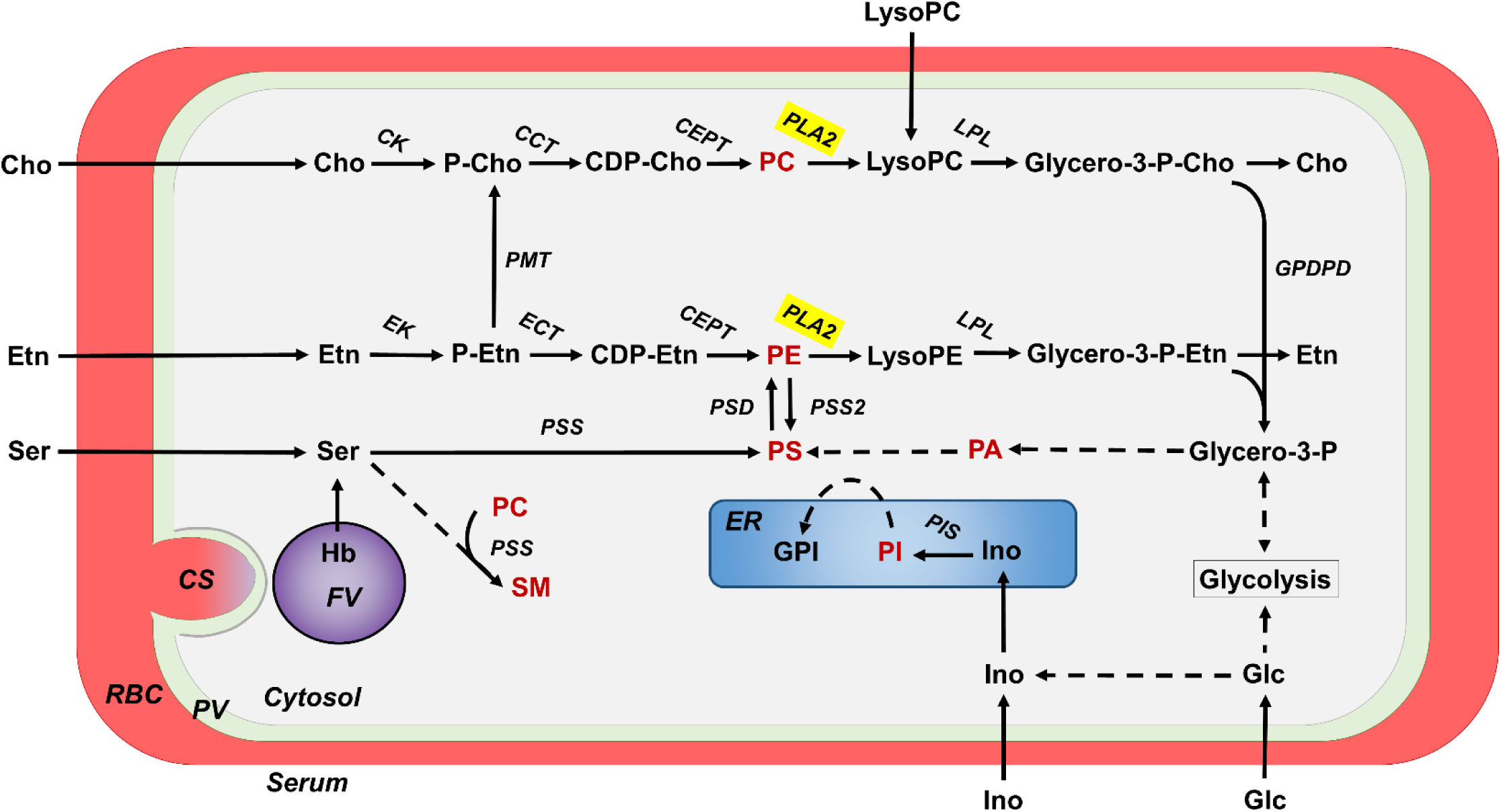
Schematic depicting the main phospholipid synthesis pathways of the *P. falciparum* blood stages. CCT, Cho-phosphate cytidyltransferase; CEPT, choline/Etn phosphotransferase; CDP-Cho, cytidine diphosphate-Cho; CDP-Etn, cytidine diphosphate ethanolamine; Cho, choline; CK, choline kinase; CS, cytostome; ECT, etanolamine-phosphate cytidyltransferase; ER, endoplasmatic reticulum; Etn, ethanolamine; FV, food vacuole; Glc, glucose; Glycero-3-P, glycerol-3-phosphate; GPDPD, glycerophosphodiester phosphodiesterase; GPI, glycosylphosphatidylinositol; Hb, hemoglobin; Ino, inositol; LPL, lysophospholipase; lysoPC, lysophosphatidylcholine; lysoPE, lysophosphatidylethanolamine; PA, phosphatidic acid; PC, phosphatidylcholine; P-Cho, phosphate-Cho; PI, phosphatidylinositol; PIS, phosphatidylinositol synthase ; PLA2, phospholipase A_2_; PMT, phosphoethanolamine N-methyltransferase; PLMT, phospholipid methyltransferase; PE, phosphatidylethanolamine; P-Etn, phosphoethanolamine; PS, phosphatidylserine; PSD, PS decarboxylase; PSS, PS synthase; PV, parasitophorous vacuole; Ser, serine; SM, sphingomyelin.

Similar to the asexual blood stages, gametocytes have a roughly 6-fold higher phospholipid level compared to the non-infected RBC. In gametocytes, however, the PC levels are lower than in the asexual blood stages (Tran et al., 2016). Under normal serum conditions, PC is generated from choline by the *de novo* cytidine diphosphate (CDP)-choline (Kennedy) pathway (reviewed in Tischer et al., 2012; Flammersfeld et al., 2018). In the absence of lysoPC in the serum, however, PC is mainly synthesized via triple methylation of phospho-ethanolamine using ethanolamine (from serum) or serine (from hemoglobin) as external precursors (Fig. 7). This pathway involves the activity of PMT and parasite lines deficient of PMT die in serum lacking these precursors (Wein et al., 2018). This alternative route of PC synthesis via the PMT pathway appears to be upregulated in gametocytes, explaining the higher tolerance of these stages towards serum-free medium (Brancucci et al., 2017).

While these data demonstrate the high sensitivity of the plasmodial blood stages to the availability of phospholipids, we now show that PNPLA1 of the parasite is involved in the phospholipid-dependent switch from asexual blood stage replication to gametocyte development. PNPLA1 is a cytosolic PLA_2_ of the asexual blood and gametocyte stages with a particular role in gametocyte induction. Asexual blood stage parasites lacking PNPLA1, due to conventional gene-KO or conditional gene-KD, are less responsive to triggers of gametocyte commitment, including lysoPC deficiency. This loss-of-function phenotype can be mimicked by the PLA_2_ inhibitor 4-BPB, pointing to direct link between PNPLA1 activity and gametocyte induction. In contrast, overexpression of PNPLA1 leads to higher gametocyte commitment rates.

The lack of PNPLA1 further results in a deregulation of the major phospholipid biosynthesis pathway in the asexual blood stages. Lipidomics indicated increased overall levels of phospholipids and more particularly PC in the absence of PNPLA1. Noteworthy, PLA_2_ activities were predicted to be important for two reactions of this pathway; the degradation of PC to lysoPC and the degradation of PE to lysoPE (see http://mpmp.huji.ac.il/). In consequence, the loss of PNPLA1 activity would evidently result in an accumulation of PC, PE, and PS, and could also lead to an increased conversion of PC into SM (Fig. 7).

PNPLA1 is one of four PNPLAs identified in *P. falciparum*, and to date for none of these functional data have been reported. However, in the Apicomplexan parasite *Toxoplasma gondii*, three of the roughly six annotated PNPLAs have meanwhile been functional characterized. TgPL1 and *Tg*PLA_2_ localize to vesicles and are discharged upon immune stress and during invasion, respectively; Furthermore, TgPL2 is crucial for apicoplast integrity (Cassaing et al., 2000; Lévêque et al., 2017; Mordue, Scott-Weathers, Tobin, & Knoll, 2007; Tobin & Knoll, 2012). These findings indicate diverse roles of the enzyme family for parasite survival. Similar diverse functions were reported for other eukaryotic PNPLAs. For example, in tobacco, PNPLAs are usually synthesized upon virus infection to generate signaling molecules that induce programmed cell death of the infected tissue (e.g. Dhondt *et al*., 2000; Dhondt *et al*., 2002; reviewed in Ryu, 2004). Mammalian PNPLAs, in contrast, are mostly used in lipid metabolism and turnover, e.g. the triglyceride synthesis and lipolysis (reviewed in Wilson and Knoll, 2018). Noteworthy, PNPLAs can also be found in prokaryotic microbes like *Legionella pneumophila*. *Legionella* injects the virulence factor VipD into the host cell, which has PLA_1/2_ activity (Gaspar & Machner, 2014; Ku et al., 2012; Shohdy, Efe, Emr, & Shuman, 2005; Zhu, Hammad, Hsu, Mao, & Luo, 2013). VipD localizes to early endosome membranes. Dependent on its PLA1 activity, it alters associated lipids and proteins and thereby avoids endosomal fusion (Gaspar & Machner, 2014). Another report described that PLA_2_-acting VipD hydrolyses PC and PE of the mitochondrial membrane, which leads to the formation of reactive lysophospholipids. These, in consequence, help to destabilize the mitochondrial membrane, resulting in the programmed killing of the infected host cell (Zhu et al., 2013).

In view of our data, we postulate that that the high PC levels due to PNPLA1-deficiency prevent malaria parasites from entering the sexual pathway. In consequence, the molecular switch from asexual blood stage replication to gametocyte formation would be flipped by decreasing PC levels. Due to reduced need of PC and the activation of the PMT pathway, gametocytes have a higher viability in serum with low levels of phospholipid precursors. Future studies need to identify, how the malaria parasite monitors its phospholipid levels and unveil the downstream signaling pathways leading to the activation of transcriptional regulators of sexual commitment.

## 4 EXPERIMENTAL PROCEDURES

### 4.1 Gene Identifiers

The following PlasmoDB gene identification numbers are assigned to the genes and proteins investigated in this study: ActinI [PlasmoDB: PF3D7_1246200]; Aldolase [PlasmoDB: PF3D7_1444800]; alpha/beta hydrolase [PlasmoDB: PF3D7_1001600]; AMA1 [PlasmoDB: PF3D7_1133400]; AP2-G [PlasmoDB: PF3D7_1222600]; ApiAP2G [PlasmoDB: PF3D7_0516800]; CRT [PF3D7_0709000]; EK [PlasmoDB: PF3D7_1124600]; Falcilysin [PlasmoDB: PF3D7_1360800]; FNPA [PlasmoDB: PF3D7_1451600]; GDV1 [PlasmoDB: PF3D7_0935400]; *Pf*39 [PlasmoDB: PF3D7_1108600]; *Pf*CCp2 [PlasmoDB: PF3D7_1455800]; *Pfs*230 [PlasmoDB: PF3D7_0209000]; *Pfs*25 [PlasmoDB: PF3D7_1031000]; PMT [PlasmoDB: PF3D7_1343000]; PNPLA1 [PlasmoDB: Pf3D7_0209100]; SAMS [PlasmoDB: PF3D7_0922200]; serine tRNA-ligase [PlasmoDB: PF3D7_0717700]; Sir2A [PlasmoDB: PF3D7_1328800].

### 4.2 Antibodies

In this study, the following primary antibodies and antisera were used: rabbit anti-HA antibody (Sigma-Aldrich); rabbit polyclonal antisera against *Pfs*230 (Biogenes), *Pfs*25 (ATCC), *Pfs*16 (Lanfrancotti, Bertuccini, Silvestrini, & Alano, 2007), AMA1 (Boes et al., 2015); mouse polyclonal antisera against falcilysin (Weißbach et al., 2017), *Pf*39 (Simon et al., 2009), actin I (Ngwa et al., 2013). The generation of polyclonal mouse antisera against PNPLA1rp1 and PNPLA1rp2 is described below.

### 4.3 Bioinformatics

For 3D modelling of PNPLA1 the I-Tasser server was used (http://zhanglab.ccmb.med.umich.edu/I-TASSER/; Yang *et al*., 2015). The 3D model was visualized using the iCn3D macromolecular structure viewer (Wang, Geer, Chappey, Kans, & Bryant, 2000). For sequence alignment of homologous proteins, the Clone Manager 9 software was used. The generation of the phylogenetic tree was done with Molecular Evolutionary Genetics Analysis software version 7.0 (Kumar, Stecher, & Tamura, 2016).

### 4.4 Parasite culture

Gametocyte producing strain *P. falciparum* NF54 was cultivated *in vitro* in RPMI 1640/ HEPES medium (Gibco) supplemented with 10% v/v heat-inactivated human serum and A^+^ erythrocytes at a 2% v/v hematocrit as described (Ifediba & Vanderberg, 1981; Trager & Jensen, 1976). Medium was completed with 50 µg/ml hypoxanthine (Sigma Aldrich) and 10 µg/ml gentamicin (Gibco) and cultures were grown in an atmosphere of 5% CO_2_, 5% O_2_ and 90% N_2_ at a constant temperature of 37°C. For cultivation of PNPLA1-KO parasites, the selection drug blasticidin (InvivoGen) was added in a final concentration of 5.4 µM; for cultivation of the PNPLA1-KD line, the selection drug WR99210 (Jacobus Pharmaceutical Company) was added in a final concentration of 2.5 nM. Human serum and erythrocyte concentrate were obtained from the Department of Transfusion Medicine, University Hospital Aachen, Germany. Donor sera and blood samples were pooled and kept anonymous. The work with human blood was approved by the Ethics commission of the RWTH University Hospital. The serum-free medium –SerM was prepared as described previously by replacing 10% v/v heat-inactivated human serum with oleic and palmitic acid (Sigma Aldrich) in a final concentration of 30 µM each, sodium bicarbonate (Sigma Aldrich) in a final concentration of 24 mM and fatty acid-free BSA (Sigma Aldrich) in a final concentration of 0.39% m/v (Brancucci et al., 2017). The –SerM/lysoPC medium was prepared by adding lysoPC (Avanti Polar Lipids) to –SerM medium in a final concentration of 20 µM; -SerM/Choline medium was prepared by adding choline (Sigma Aldrich) to –SerM medium in a final concentration of 300 µM and glucose in a final concentration of 11.1 mM (Brancucci et al., 2017). To synchronize the asexual parasite blood stages, parasites cultures with 3-4% ring stages were centrifuged, the pellet was resuspended in five time pellet’s volume of 5% sorbitol (AppliChem)/ddH20 and incubated for 10 min at room temperature (RT) (Lambros & Vanderberg, 1979). Cells were washed once with RPMI to remove sorbitol and further cultivated as described above. To synchronize cultures to a 4-h time window consecutive sorbitol treatments were applied as described (Brancucci et al., 2017; Brancucci, Goldowitz, Buchholz, Werling, & Marti, 2015). Mature schizonts and gametocytes were enriched via Percoll gradient centrifugation (GE Healthcare Life Sciences) as described previously (Kariuki et al., 1998; Radfar et al., 2009). Gametogenesis was induced *in vitro* by incubating mature gametocyte cultures in 100 µM xanthurenic acid (XA) dissolved in 1% v/v 0.5 M NH_4_OH/ddH_2_0 for 15 min at RT (Billker et al., 1998; Garcia, Wirtz, Barr, Woolfitt, & Rosenberg, 1998).

### 4.5 Recombinant protein expression

Recombinant proteins corresponding to PNPLA1rp1 (spanning aa 498-670) and PNPLA1rp2 (spanning aa 335-496) were expressed as fusion proteins with an N-terminal maltose binding protein (MBP)-tag using the pMAL^TM^c5x-vector (New England Biolabs). Cloning was mediated by the addition of the restriction sites XmnI/PstI to the ends of gene fragments PCR-amplified from *P. falciparum* gDNA, using PNPLA1rp1 forward primer and PNPLA1rp1 reverse primer for PNPLA1rp1 and PNPLA1 rp2 forward primer and PNPLA1 rp2 reverse primer for PNPLA1rp2 (for primer sequences, see Table S1). Recombinant proteins were expressed using *E.coli* BL21 (DE3) RIL according to the manufacturer’s protocol (Stratagene). The recombinant fusion proteins were purified via affinity chromatography from bacterial extracts using amylose resin (New England Biolabs) according to manufacturer’s protocol. The full-length recombinant protein corresponding to PNPLA1rp3 (spanning aa 001-679) was expressed as fusion protein with an N-terminal glutathione-S-transferase (GST)-tag using the pGEX-6P-1 vector (GE Healthcare). A synthetic PNPLA1 gene, codon-optimized for recombinant protein expression in *E. coli* with the GeneOptimizerTM software (Thermo Fisher Scientific), was used as template DNA (for synthetic sequence, see Fig. S13). Cloning was mediated by the addition of the restriction sites BamHI and XhoI to the ends of gene fragments PCR-amplified from codon-optimized DNA-template using the PNPLA1rp3 forward primer and PNPLA1rp3 reverse primer (for primer sequences, see Table S1). Recombinant proteins were expressed using *E.coli* BL21 (DE3) RIL according to the manufacturer’s protocol.

### 4.6 Generation of mouse antisera

Recombinant fusion proteins PNPLA1rp1-MBP and PNPLA1rp2-MBP were purified via affinity chromatography as described above followed by PBS buffer exchange via filter centrifugation using Amicon Ultra 15 (Sigma-Aldrich) according to the manufacturer’s protocol. Protein concentrations were determined via Bradford assay. Immune sera were generated by immunization of 6 weeks-old female NMRI mice (Charles Liver Laboratories) via subcutaneous injection of 100 µg recombinant protein emulsified in Freund’s incomplete adjuvant (Sigma-Aldrich) followed by a boost after 4 weeks with 50 µg of recombinant protein. Mice were anesthetized 10 days after the boost by intraperitoneal injection of ketamine-xylazine mixture according to the manufacturer’s protocol (Sigma-Aldrich). Polyclonal immune sera were collected via heart puncture and pooled from three mice immunized with the same antigen. NMS were collected for negative control in the experiments. Experiments for the generation of antisera in mice were approved by the animal welfare committees of the District Council of Cologne, Germany (ref. no. 84-02.05.30.12.097 TVA)

### 4.7 Generation of the PNPLA1-KO parasite lines

PNPLA1-KO parasite lines were generated via single cross-over homologous recombination using the pCAM-BSD-KO vector (Bennink et al., 2018; Dorin-Semblat et al., 2007; Wirth, Bennink, Scheuermayer, Fischer, & Pradel, 2015; Wirth et al., 2014). A 567 bp gene fragment homologous to the PNPLA1 gene coding for the N-terminal part was amplified via PCR using the PNPLA1-KO forward and reverse primers (for primer sequences, see Table S1). Ligation of insert and vector backbone was mediated by BamHI/ NotI restriction sites. A WT culture synchronized for 5% ring stages was loaded with 100 µg vector in transfection buffer via electroporation (310 V, 950 µF, 10 ms; Bio-Rad gene-pulser) as described (Ngwa et al., 2017; Wirth et al., 2014). Blasticidin (InvivoGen) was added to a final concentration of 5.4 µM at 6h post-transfection. A mock control was electroporated using transfection buffer only without addition of vector-DNA und cultured in regular A+-medium. Blasticidin-resistant parasites appeared after 3-4 weeks. To confirm successful plasmid integration into the *pnpla1* gene locus, genomic DNA (gDNA) of transfected parasites was isolated using the NucleoSpin Blood Kit (MACHEREY-NAGEL) according to the manufacture’s protocol and used as template in diagnostic PCR. The following primers were used to confirm correct integration of the vector into the target locus: PNPLA1-KO 5’-integration forward primer (1), PNPLA1-KO 3’-integration reverse primer (2), pCAM-BSD forward primer (3) and pCAM-BSD reverse primer (for primer location see Fig. S4; for primer sequences, see Table S1). Successful confirmation of vector integration was followed by clonal dilution of a >3% ring stage culture in a 96-well plate. After three weeks of cultivation, clonal sub-cultures were identified via Malstat assay as described below and subsequent diagnostic PCR was performed to confirm correct integration and absence of the WT *pnpla1* gene locus. Two clonal lines, PNPLA1-KO A12 and PNPLA1-KO C11, were isolated.

### 4.8 Generation of the PNPLA1-KD parasite line

PNPLA1-KD parasite lines were generated via single cross-over homologous recombination using the pARL-HA-*glmS* vector (Fig. S3). An 867-bp gene fragment homologous to the PNPLA1 gene coding for the C-terminal part was amplified using the PNPLA1-KD forward primer and the PLPLA1-KD reverse primer (for primer sequences, see Table S1). The stop codon was excluded from the homologous gene fragment. Ligation of the insert with the vector backbone was mediated by NotI and AvrII restriction sites. Transfection of parasites was performed as described above. For selection of parasites carrying the vector, WR92210 (Jacobus Pharmaceutical Company) was added to a final concentration of 2.5 nM and successful integration of the vector was confirmed by diagnostic integration PCR using PNPLA1-KD 5’-integration primer (1), PNPLA1-KD 3’-integration primer (2), pARL-HA-*glmS* forward primer (3) and pARL-HA-*glmS* reverse primer (4) (for primer location see Fig. S6; for primer sequences, see Table S1).

### 4.9 Generation of the PNPLA1-GFP episomal expression parasite lines

The *crt*-PNPLA1-GFP and the *fnpa*-PNPLA1-GFP parasite lines were generated using the pARLII-GFP vector (Fig. S10A). The *pnpla1* full-length gene was amplified from cDNA using the PNPLA1-gfp forward primer and the PNPLA1-gfp reverse primer (for primer sequences, see Table S1). The stop codon was excluded from the full-length gene sequence. Ligation of the insert with the vector backbones was mediated by XhoI and AvrII. The plasmid was sequenced to confirm that the encoding segment was inserted in frame with the GFP encoding sequence. Transfection of parasites was performed as described above. For selection of parasites carrying the vectors, WR92210 (Jacobus Pharmaceutical Company) was added to a final concentration of 2.5 nM and successful uptake of the vector was confirmed by diagnostic PCR using the PNPLA1-gfp episome forward primer and the PNPLA1-gfp episome reverse primer (for primer location see Fig. S10; for primer sequences, see Table S1).

### 4.10 Indirect immunofluorescence assay

Mixed asexual blood stage cultures, mixed gametocyte cultures, macrogametes as well as zygotes, collected at 15 min, or 30 min, of the WT, the PNPLA1-KD and the PNPLA1-KO lines A12 and C11 were air-dried as cell monolayers on glass slides and subsequently fixed with 4% w/v paraformaldehyde/PBS (pH 7.4) for 10 min at RT. For membrane permeabilization, the fixed parasites were incubated with 0.1% v/v Triton X 100/125 mM glycerine for 10 min, followed by blocking of non-specific binding sites using 3% w/v BSA/PBS for 30 min at RT. The preparations were incubated with polyclonal mouse antisera specific for PNPLA1rp1 and PNPLA1rp2 (dilution 1:200), with rabbit anti-HA antiserum (dilution 1:200) or with NMS (dilution 1:200) for 2 h at 37°C. Binding of the primary antibody was detected by incubation with Alexa Fluor 488-conjugated goat anti-mouse or secondary antibody (Invitrogen Molecular Probes). The sexual stages were highlighted by double-labelling with rabbit antibodies directed against *Pfs*230, followed by incubation with polyclonal Alexa Fluor 594-conjugated goat anti-rabbit secondary antibodies (Invitrogen Molecular Probes), while the asexual blood stages were stained with 0.001% w/v Evans Blue (EB) (Sigma Aldrich)/PBS for 3 min at RT followed by 5 min washing with PBS. The parasite nuclei were highlighted by treatment with Hoechst33342 nuclear stain (Invitrogen) for 1 min at RT. Cells were washed with PBS and mounted with anti-fading solution AF2 (CitiFluor^TM^) and sealed with nail polish. Specimen were examined with a Leica DM 5500 B microscope and digital images were processed using the Adobe Photoshop CS software.

### 4.11 Merozoite quantification

Cell monolayers of synchronized mature schizonts of WT and of PNPLA-KO A12 and PNPLA1-KO C11 were air-dried on glass slides and subsequently fixed with 4% w/v paraformaldehyde/PBS (pH 7.4) for 10 min. Counterstaining with EB and highlighting of nuclei by Hoechst33342 nuclear stain was performed as described above. Specimen were examined with a Leica DM 5500 B microscope and number of nuclei of mono-nucleated merozoites were counted for 75 schizonts per setting. Only mature schizonts that had clearly segmented merozoites with limited overlap and with single pigmented residual bodies indicating singular infections were used for the analysis.

### 4.12 Diagnostic RT-PCR

Total RNA was isolated from sorbitol-synchronized ring, trophozoite and schizont cultures, as well as from Percoll-gradient enriched immature (stage II-IV), mature (stage V) and activated gametocytes (15 min p.a.) of the WT using the Trizol reagent (Invitrogen) according to the manufacture’s protocol. RNA was prepared by phenol/chloroform preparations followed by ethanol precipitation and subsequent treatment with RNase-free DNase I (Qiagen). Photometric analyses of RNA samples all had A260/280 ratios higher than 2.1. Two µg of each RNA sample were used as template for cDNA synthesis using the SuperscriptIV First-Strand Synthesis System (Invitrogen) according to manufacturer’s protocol. Transcript for PNPLA1 (220 bp) was amplified in 25 cycles using PNPLA1 RT forward and reverse primers (for primer sequences, see Table S1). To confirm purity of the asexual blood stage and gametocyte samples, transcript amplifications of *ama1* (407 bp) using AMA1-RT forward and reverse primers and of *pfccp2* (286 bp) using CCP2-RT forward primer and reverse primers were performed (Peterson et al., 1989; Pradel et al., 2004). Amplifications of the housekeeping gene *aldolase* (378 bp) using Aldolase-RT forward and reverse primers were used as loading control and to test for residual genomic DNA in the negative controls without reverse transcriptase. PCR products were separated by 1.5% agarose gel electrophoresis.

### 4.13 Real-time RT-PCR

RNA from asexual blood stages was isolated as described above and 2 µg of each total RNA sample was used as template for cDNA synthesis using the Super Script IV First-Strand Synthesis System (Invitrogen) following the manufacturer’s protocol. The synthesized cDNA was first tested by diagnostic RT-PCR for purity of the asexual blood stage and gametocyte samples, using *ama1-* and *pfccp2-*specific primers, respectively. Controls without reverse transcriptase were used to exclude potential genomic DNA contamination by using *aldolase* primers. Primers for real-time PCR were designed using the Primer 3 software (Primer3 Input; version 0.4.0) and were initially tested for specificity via conventional PCR on gDNA. Real-time RT-PCR measurements were performed using the StepOnePlus Real-Time PCR System (Applied Biosystems). Samples were analyzed in triplicate in a total volume of 20 µl using the maxima SYBR green qPCR master mix according to manufacturer’s instructions (Thermo Scientific). Controls without template and without reverse transcriptase were included in all real-time RT-PCR experiments. Transcript expression levels were calculated by the 2^-ΔΔCt^ (Livak & Schmittgen, 2001) method using the endogenous control gene encoding the *P. falciparum* seryl tRNA-ligase (PF3D7_0717700) as reference (Bennink et al., 2018; Ngwa et al., 2017; Salanti et al., 2003).

### 4.14 Subcellular fractioning

The protocol for subcellular fractionation of parasite lysates using the solubility of proteins in different buffers was modified from Grüring et al. (2012). In brief, ∼3*10^6 schizonts of the PNPLA1-KD line were purified via Percoll gradient and liberated from the host cells with 0.03% saponin/PBS for 3 min at 4°C. After washing with PBS, parasites were hypotonically lysed in 100 µl of 5 mM Tris-HCL (pH 8), supplemented with protease inhibitor cocktail (complete EDTA-free, Roche) for 10 min at RT followed by freezing at −20°C and thawing on ice. Soluble proteins (Tris-HCL fraction) were separated from non-soluble constituents by centrifugation (5 min, 16,000xg, 4°C). This was followed by sequential pellet extraction with 100 µl each of freshly prepared Na_2_CO_3_ on ice for 30 min (peripheral fraction), ice-cold 1% Triton X-100 (integral membrane fraction), and 0.5 x PBS/4% SDS/ 0.5% Triton X-100 (insoluble fraction) at RT. Between each extraction, the remaining pellet was washed with 1 ml ice-cold PBS to remove any impurities. Supernatants were centrifuged to remove residual material. The individual fractions were subjected to WB as described below.

### 4.15 Western blotting

Asexual blood stage parasites of the WT, the PNPLA1-KO lines A12 and C11, and the PNPLA1-KD line were harvested from mixed or synchronized cultures, while gametocytes were enriched by Percoll purification. Parasites were released from iRBCs with 0.015% w/v saponin/PBS for 10 min at 4°C, washed with PBS and resuspended in lysis buffer (0.5% Triton X-100, 4% w/v SDS, 0.5xPBS) supplemented with protease inhibitor cocktail (Roche). 5xSDS-PAGE loading buffer containing 25 mM dithiothreitol was added to the lysates, samples were heat-denatured for 10 min at 95°C, and separated via SDS-PAGE. Following gel electrophoresis, separated parasite proteins were transferred to Hybond ECL nitrocellulose membrane (Amersham Biosciences) according to the manufacturer’s protocol. Non-specific binding was blocked by incubation of the membranes in Tris-buffered saline containing 5% w/v skim milk, pH 7.5, followed by immune recognition overnight at 4°C using polyclonal mouse immune sera specific for *Pf*39 (dilution 1:5000), PfAMA1 (dilution 1:1,000), falcilysin (1:1,000), actin I (1:1,000), PNPLA1rp1 (1:1,000), PNPLA1rp2 (1:1,000) or using polyclonal rabbit immune sera specific to anti-HA (1:1,000). After washing, membranes were incubated with the respective alkaline phosphatase-conjugated secondary antibody (Sigma-Aldrich) for 1 h at RT and developed in a solution of nitroblue tetrazolium chloride (NBT) and 5-brom-4-chlor-3-indoxylphosphate (BCIP; Sigma-Aldrich) for 5-30 min at RT. Blots were scanned and processed using the Adobe Photoshop CS software. Band intensities were measured using the ImageJ programme version 1.51f.

### 4.16 Asexual blood stage replication assay

To compare asexual blood stage replication between the parental WT and the PNPLA1-KO lines A12 and C11, synchronized asexual blood stage cultures were set to an initial parasitemia of 0.25% ring stages und cultivated as described above. Giemsa-stained thin blood smears were prepared every 24 h over a time-period of 120 h at six different time points (0, 24, 48, 72, 96, 120 h post-seeding). Parasitemia of each time point was determined microscopically at 1,000-fold magnification by counting the percentage of parasites in 1,000 RBCs. To identify the blood stages (ring, trophozoites, schizonts, mature schizonts) present in the cultures at a given time point, 100 iRBCs were counted per setting. To analyze asexual blood stage replication and length of the intraerythrocytic replication cycle in the PNPLA1-KD parasite line, experiments were set-up as described above. Expression of PNPLA1 was knocked-down in transgenic PNPLA1-KD parasite line by treatment of the culture with GlcN (D-(+)-glucosamine hydrochloride; Sigma Aldrich) at a final concentration of 2.5 mM. GlcN treatment started one cycle before the experimental set-up. The GlcN-treated culture was compared to the untreated culture cultivated in normal cell culture medium. To exclude an unspecific effect of GlcN, the parental WT was used as control. For each assay, three experiments were performed, each in triplicate. Data analysis was performed using MS Excel 2014.

### 4.17 Gametocyte development assay

#### PNPLA1-KO lines

To compare gametocyte development between the WT and the PNPLA1-KO lines A12 and C11, synchronized ring stage cultures were set to a parasitemia of 2% with 5% hematocrit and cultivated in cell culture medium at 37°C. The medium was exchanged daily. Cultures were either maintained in cell culture medium over a time period of 14 d and samples were taken in triplicate every 24 h, or, when cultures reached a parasitemia of 10%, gametocytogenesis was induced by addition of lysed RBCs for 24 h (Ngwa et al., 2017). Cells were washed and cultures were maintained in cell culture medium supplemented with 50 mM N-acetyl glucosamine (GlcNAc) for ∼5 d to kill the asexual blood stages followed by cultivation in normal cell culture medium (Fivelman et al., 2007; Ngwa et al., 2017). Samples were taken in triplicate every 24 h starting from day 2 p.g.i. for Giemsa smear preparation. To induce sexual differentiation with –SerM medium, tightly synchronized parasites (28 ± 2 h.p.i.) with a 1% parasitemia at 5% hematocrit were incubated for 22 h in –SerM as described (Brancucci et al., 2017). Incubation with –SerM was followed by cultivation in cell culture medium supplemented with 50 mM GlcNac to kill the asexual blood stages. Samples were taken in triplicate every 24 h starting from day 2 p.g.i. for Giemsa smear preparation. Cultures cultivated in –SerM/lysoPC, –SerM/Choline and –SerM/Serum were used as control (Brancucci et al., 2017).

#### PNPLA1-KD line

To analyze gametocytogenesis in the PNPLA1-KD line, experiments were set-up as described above. Expression of PNPLA1 was knocked-down in the PNPLA1-KD parasite line by treatment of the culture with the inducer GlcN (Sigma Aldrich) at a final concentration of 2.5 mM. GlcN was added one cycle before the experimental set-up. The GlcN-treated culture was compared to the untreated cultures. To exclude an unspecific effect of GlcN, the parental WT was used as control. Gametocytemia was determined per 1,000 RBCs and the gametocyte stages II-IV at the different time points were recorded. For each assay, three experiments were performed, each in triplicate. Data analysis was performed using MS Excel 2014 and GraphPad Prism 5.

### 4.18 Comparative exflagellation assay

At day 14 p.g.i., mature gametocytes of the WT and the PNPLA1-KO lines A12 and C11 were adjusted with erythrocytes to a final gametocytemia of 1% with a total hematocrit of 5% in cell culture medium. A volume of 100 µl of each gametocyte culture was activated *in vitro* with 100 µM XA for 15 min at RT. At 15 min p.a., the numbers of exflagellation centers were counted at 400-fold magnification in 30 optical fields using a Leica DMLS microscope. Four independent experiments were conducted in quadruplicate; data analysis was performed using MS Excel 2014. Exflagellation was calculated as a percentage of the number of exflagellation centers in the PNPLA1-KO lines in relation to the number of exflagellation centers in the WT control (WT set to 100%).

### 4.19 Comparative macrogamete and zygote formation assay

For the analysis of macrogamete and zygote formation, mature gametocytes of the WT and of the PNPLA1-KO lines A12 and C11 were enriched via Percoll purification and adjusted to a final gametocytemia of 2% with a total hematocrit of 5% in cell culture medium. Gametocyte cultures were activated *in vitro* with 100 µM XA for 30 min (macrogametes) or 6 h (zygotes) at RT. An equal volume of each sample was coated on Teflon slides and subjected to IFA as described above using rabbit anti-*Pfs*25 antisera for macrogamete and zygote labelling. Parasites were counted microscopically using a Leica DM 5500B fluorescence microscope with 600-fold magnification. Numbers of macrogametes and zygotes were counted for a total number of 1000 erythrocytes in triplicate and calculated in relation to the WT control (set to 100%). For each assay, three experiments were performed, each in triplicate. Data analysis was performed using MS Excel 2014.

### 4.20 Malstat assay

To determine the antimalarial effect of the phospholipase inhibitors 2,4’-dibromoacetophenone (4-BPB; Sigma Aldrich), ASB14780 (Sigma Aldrich), N-(p-amylcinnamoyl) anthranilic acid (Biomol), and bromoenol lactone (Biomol), a Malstat assay was performed as described previously (Aminake et al., 2011). Synchronized ring stages of the WT were plated in triplicate in a 96-well plate (200 µl/ well) at a parasitemia of 1.0% in the presence of the respective inhibitor dissolved in DMSO (500 µM to 0.05 nM for 4-BPB and N-(p-amylcinnamoyl) anthranilic acid and 1mM to 7.8 µM for bromoenol lactone and ASB14780). Chloroquine, dissolved in RPMI, served as positive control, DMSO alone at a concentration of 0.5% v/v was used as negative control. Parasites were cultivated *in vitro* in an airtight container as described above. After 72 h, 20 µl of each well were mixed with 100 µl Malstat reagent in a new 96-well microtiter plate. Parasite lactate dehydrogenase activity was measured by adding 20 µl of a 1:1 mixture of 1 mg/ml diaphorase and 1 mg/ml NBT (nitroblue tetrazolium) to the Malstat reaction, and optical densities were measured at 630 nm. Three experiments were performed, each in triplicate. The IC_50_ values were calculated from variable-slope sigmoidal dose-response curves using the GraphPad Prism 5 program.

### 4.21 Gametocyte toxicity assay

WT parasites were grown at high parasitemia to induce gametocyte formation. When stage II gametocytes were formed, 1 ml of the culture was plated in triplicate in 24-well plate and incubated with 4-BPB at IC_50_ (5.62 µM) concentration. Chloroquine at IC_50_ (12.1 nM) concentration and 0.5% v/v DMSO served as negative controls. The proteasome inhibitor epoxomicin (60 nM) was diluted in DMSO and used as positive control (Aminake et al., 2011; Ngwa et al., 2017). The parasites were treated with inhibitors for 2 d and the medium was replaced daily. The parasites were cultivated for 10 d and Giemsa-stained blood smears were taken at day 7 and day 10 to determine numbers of gametocytes of stages IV and V per 1,000 RBCs. Three experiments were performed, each in triplicate. Data analysis was performed using MS Excel 2014 and GraphPad Prism 5.

### 4.22 Lipidomics analysis

Analysis was performed on four independent cell harvests of the WT and the PNPLA1-KO lines A12 and C11. Highly synchronous parasite cultures were harvested by transferring the cultures to pre-warmed 50-ml tubes and the total amount of parasites were determined by Giemsa staining and via counting on a haemocytometer. The cultures were metabolically quenched via rapid cooling to 0°C by suspending the tube over a dry ice/100% ethanol slurry mix, while continually stirring the solution. Following quenching, parasites were constantly kept at 4°C and liberated from the enveloping RBCs by mild saponin treatment with 0.15% w/v saponin/0.1% w/v BSA PBS for 5 min. After three washing steps with 1 ml ice-cold PBS followed by centrifugation each time, the parasite pellet was subjected to lipid extraction using chloroform and methanol (Amiar et al., 2016) in the presence of butylhydoxytluene, PC (C21:0/C21:0), C13:0 fatty acids and Lipidomix lipid standard mix (Avanti). Then the extract was set to biphase separation with 0.2% KCl to remove polar molecules. The total lipid extract was dried under the nitrogen gas. The lipid was then dissolved in the 1-butanol and separated by two dimension high performance thin layer chromatography (NanoSiL, MACHEREY-NAGEL). For the first dimension, the solvent system of chloroform/methanol/28% ammonium hydroixide (65:35:8, v/v/v) and for the second dimension, chloroform/acetone/methanol/acetic acid/water, 40:15:14:13:7.5 (v/v/v/v/v) were used, respectively. Each lipid spot was confirmed with authentic standards. The lipid spot was then scraped off and set to methanolysis with C15:0 fatty acid standard in 0.5 M HCl/ methanol at 85°C for 6 h. Generated fatty acid methyl esters were then extracted by hexane. Fatty acid methyl esters were detected by GC-MS (Agilent 5977A-7890B) according to the method described in Ramakrishnan et al. (2012) and quantified by Mass Hunter software (Agilent). Each fatty acid was quantified according the calibration curve generated with authentic fatty methyl ester standards.

### 4.23 Statistical analysis

Data are expressed as mean ± SD. Statistical differences were determined using One-Way ANOVA with Post-Hoc Bonferroni Multiple Comparison test or unpaired tow-tailed Student’s t-test, as indicated. P-values <0.05 were considered statistically significant. Significances were calculated using GraphPad Prism 5 and are represented in the figures as follows: ns, not significant p>0.05; * p<0.05; **p<0.01; ***p<0.001.

## Acknowledgements

The authors thank Matthias Marti (University of Glasgow) and Chetan Chitnis (Pasteur Institute Paris) for helpful discussions. The authors further thank Christina Lang (Robert Koch Institute) and Alexandra Golzmann and Sofia Basova (RWTH Aachen University) for support with the experiments. The work was funded by the priority programme SPP1580 of the Deutsche Forschungsgemeinschaft (GP, AF, and JMP).

---

**TABLE S1.**
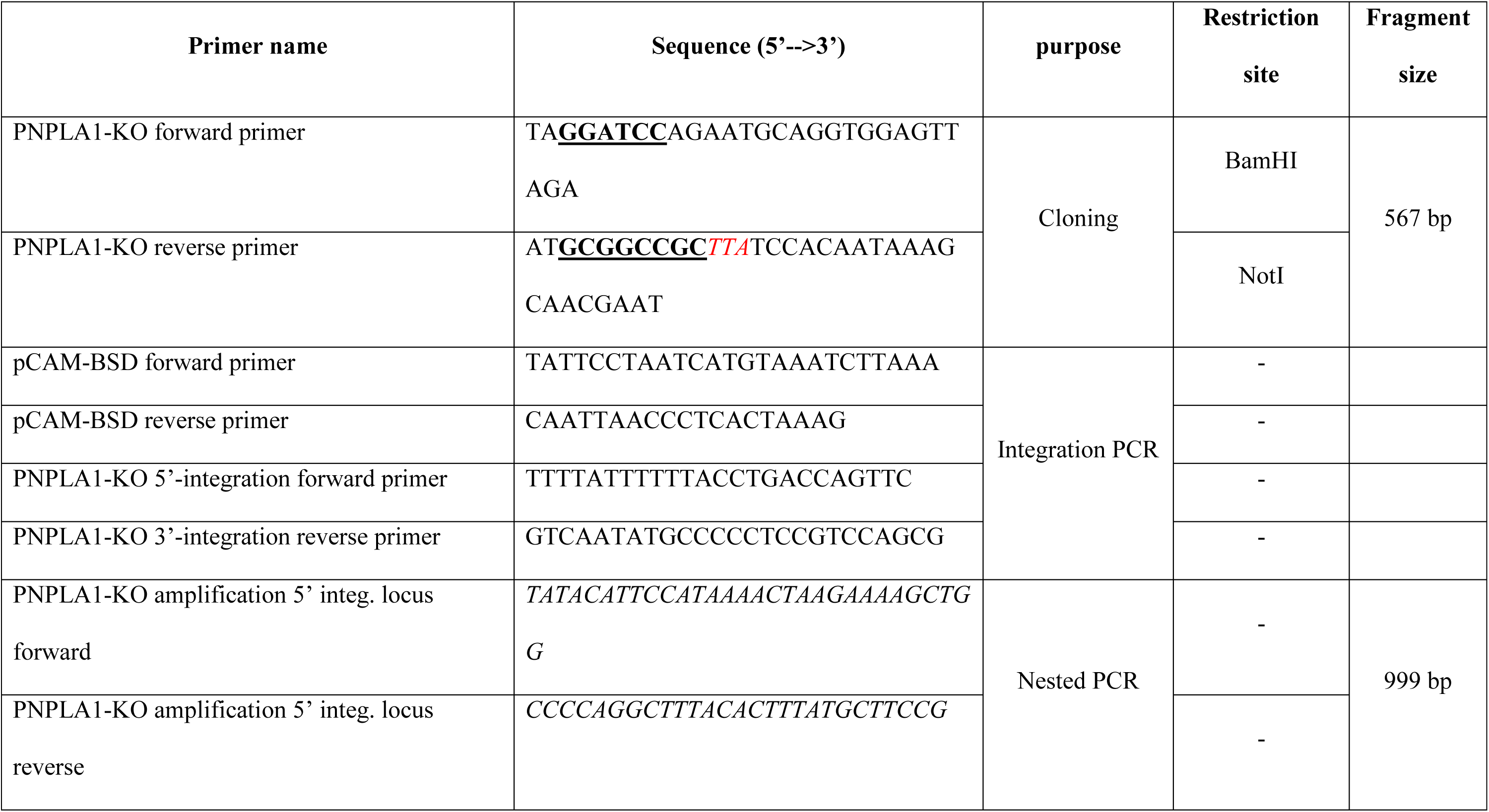
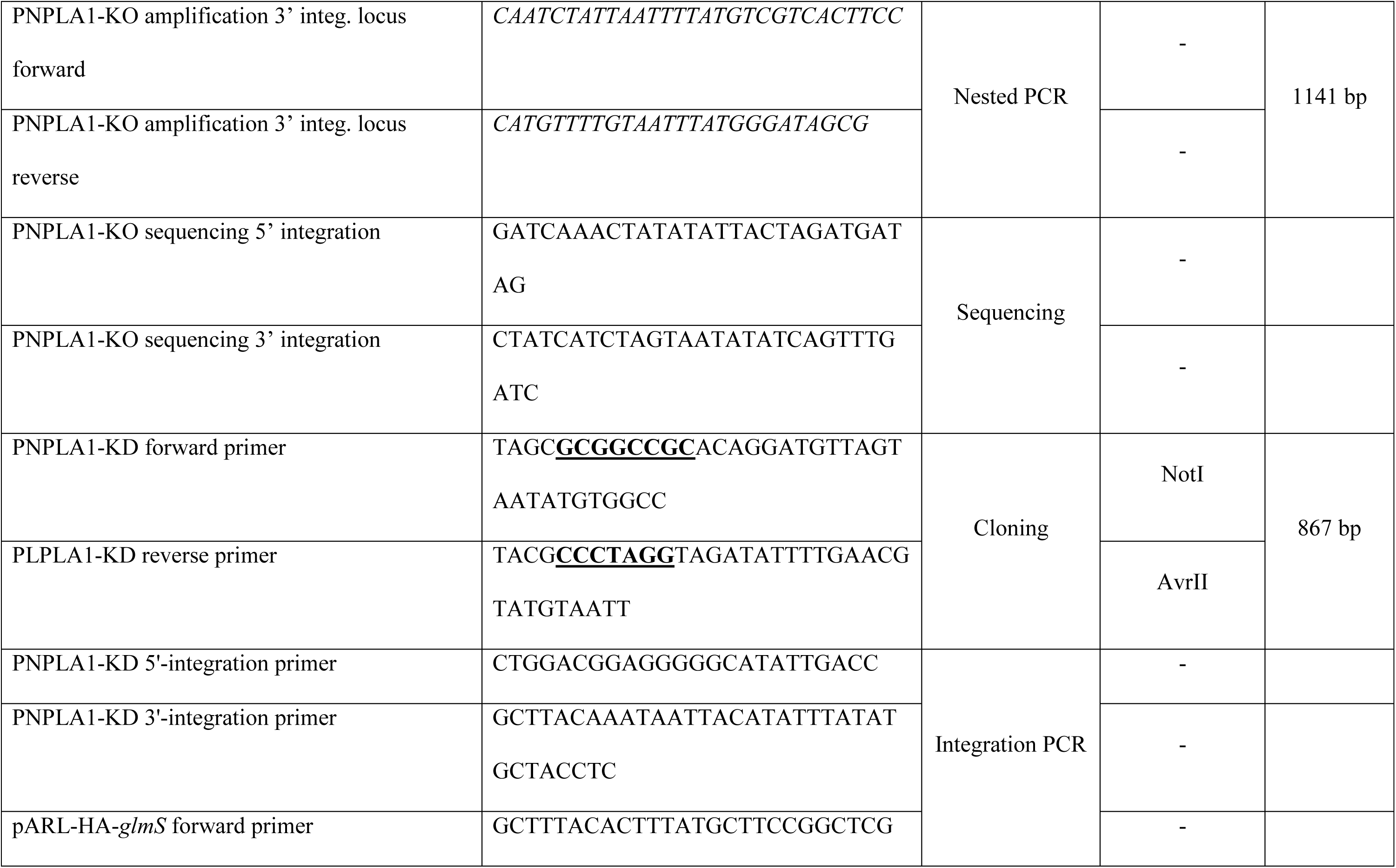
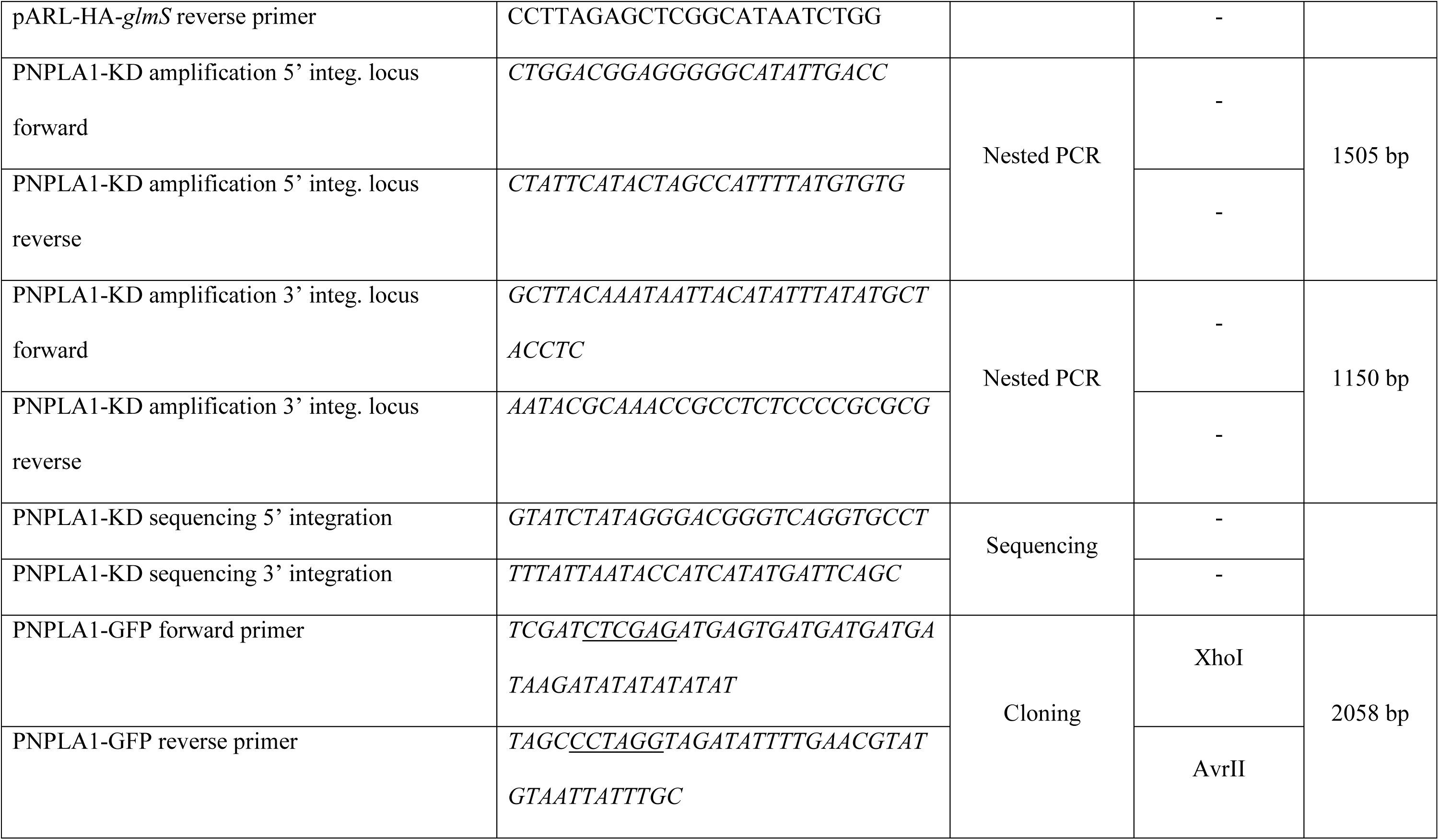
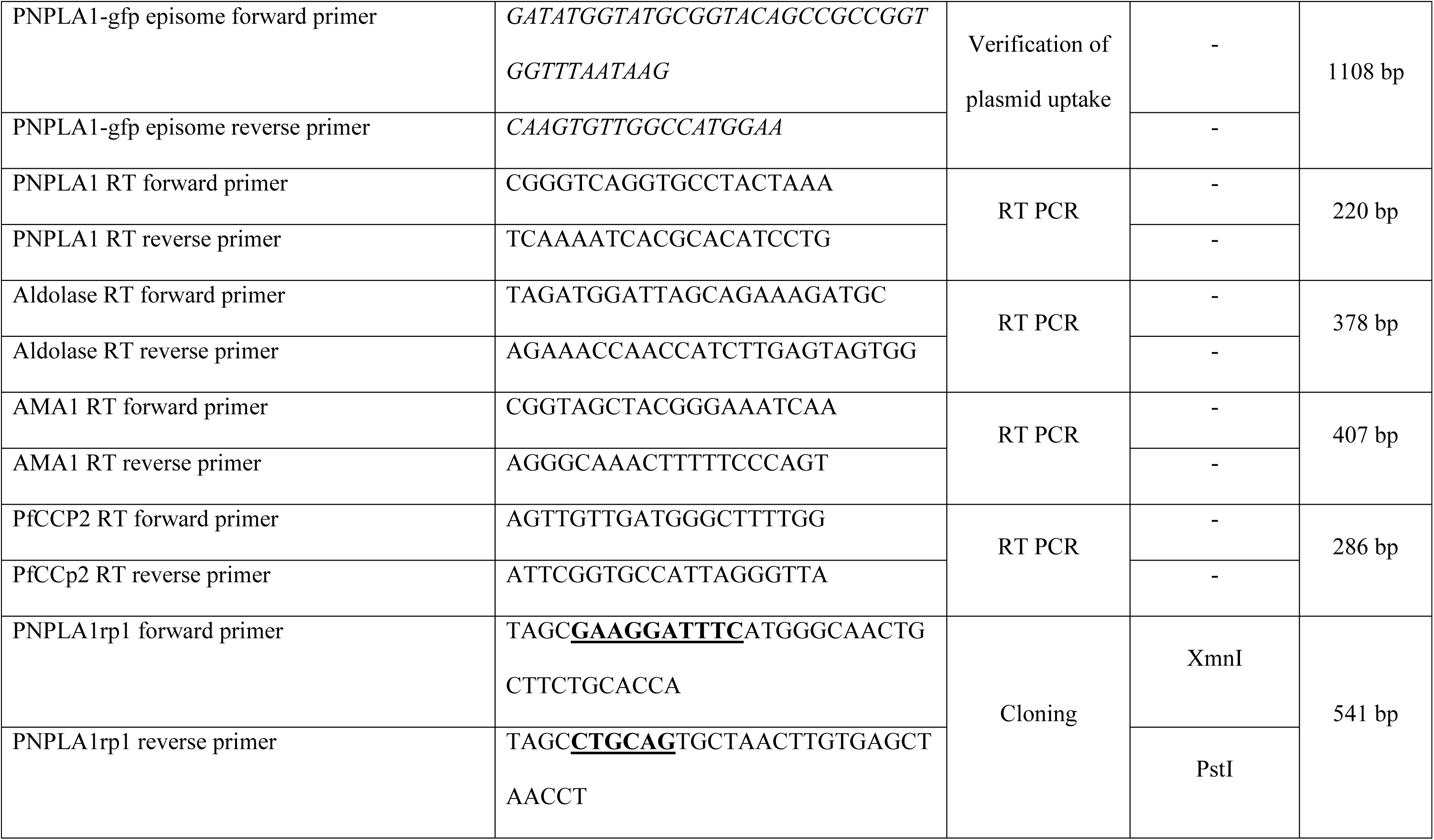
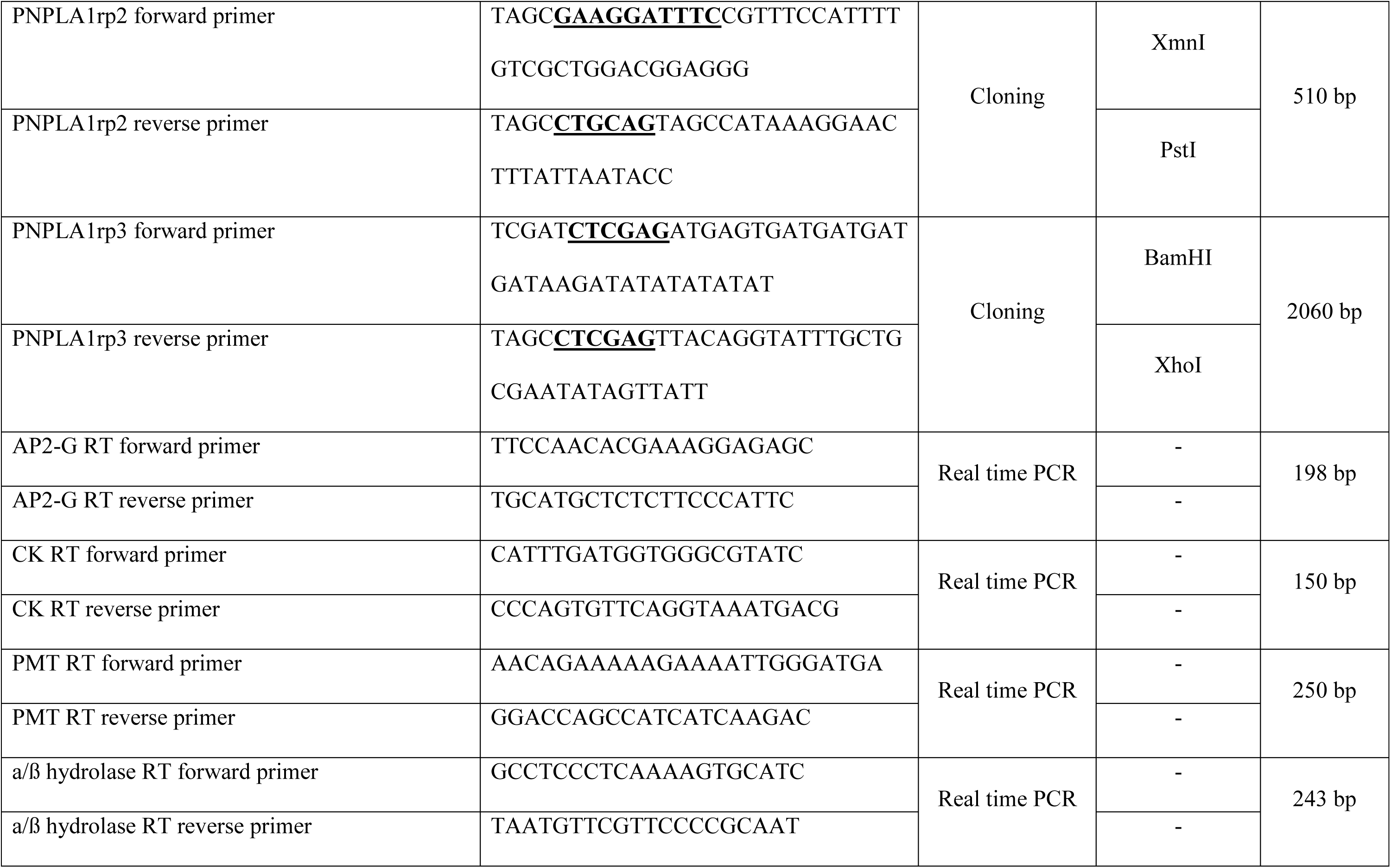
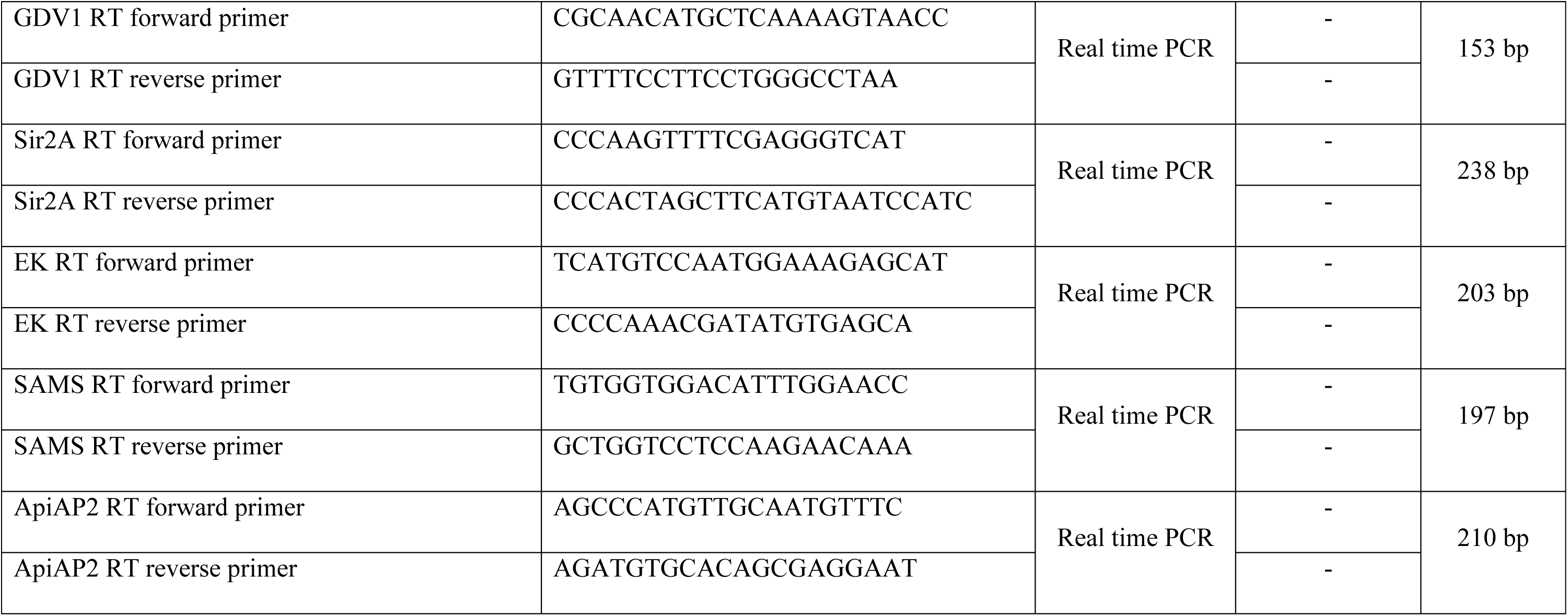
List of primers used in this study.

## SUPPLEMENTAL FIGURE LEGENDS

**FIGURE S1.**
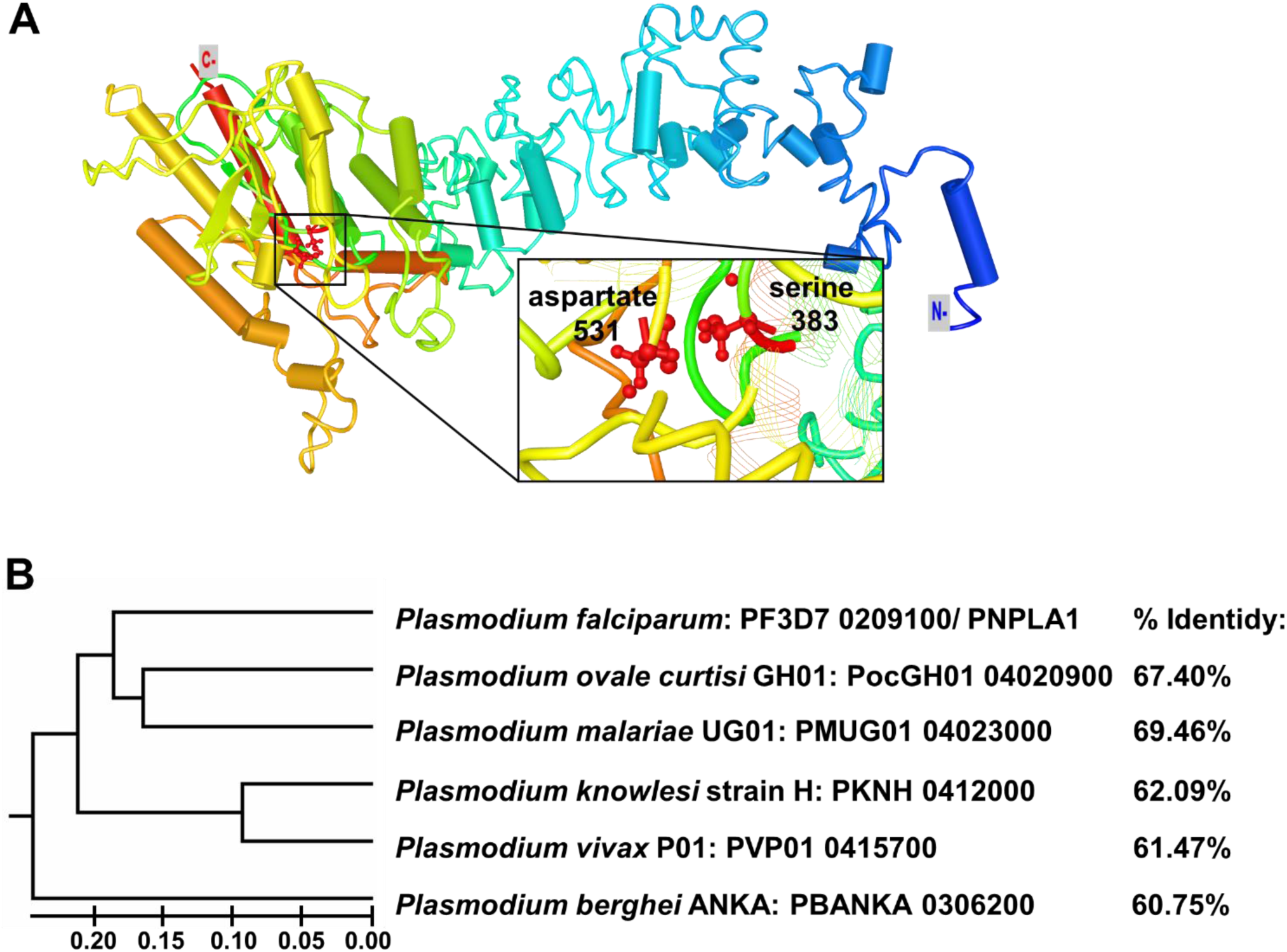
3D structure and phylogenetic analysis of PNPLA1. **(A)** The 3D structure of PNPLA1. The architecture was modelled using the I-TASSER server based on the amino acid sequence and the crystal structure of PDB hit 6AUN (C-score −1.61, identity 0.156, coverage 0.842), a calcium-independent phospholipase A_2_-β. The sequence is shown from the N-terminus (blue) to the C-terminus (red). The putative catalytic dyad, consisting of the coordinated serine and aspartate, are highlighted. **(B)** Phylogenetic analysis of PNPLA1 of *P. falciparum* and orthologous proteins in other *Plasmodium* species. Phylogenetic analysis was performed using MEGA7. Percentage of identities are indicated and were determined with the NCBI Basic Local Alignment Search Tool (BLAST).

**FIGURE S2.**
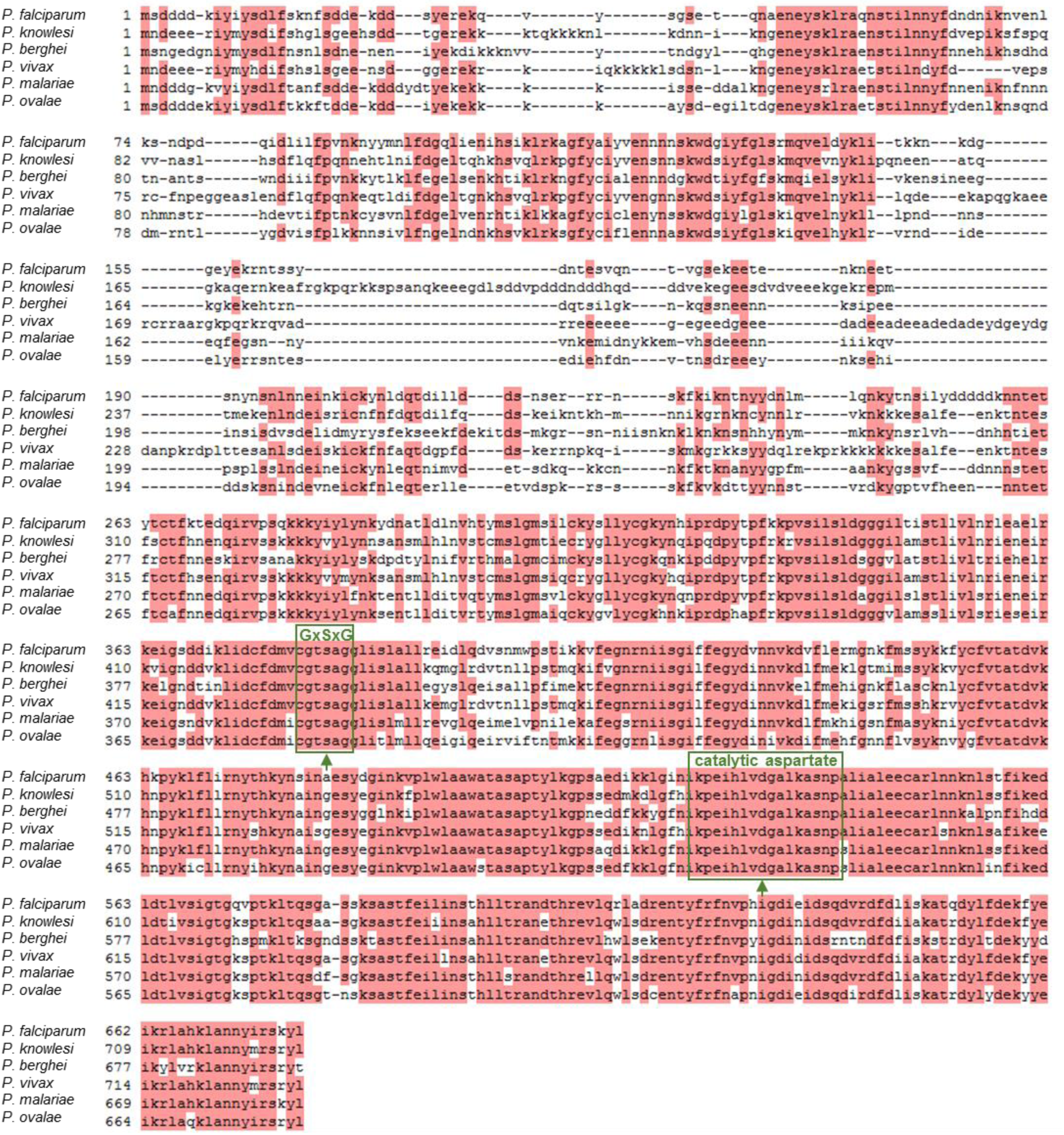
Sequence alignment of PNPLA1. Amino acid sequence alignment of PNPLA1 and related proteins in other *Plasmodium* species identified with the NCBI Local Alignment Search Tool (BLAST). Regions with sequence homology ≥ 60% are highlighted in red. The putative catalytic serine that is embedded in the Gly-Xaa-Ser-Xaa motif and the region comprising the putative catalytic aspartate are framed in green. Sequence alignment was generated with the Clone Manager 9 Software. *Plasmodium falciparum*, PF3D7_0209100; *Plasmodium ovale*, PocGH01_04020900; *Plasmodium malariae*, PMUG01_04023000; *Plasmodium knowlesi*, PKNH_0412000; *Plasmodium vivax*, PVP01_0415700*; Plasmodium berghei*, PBANKA_0306200.

**FIGURE S3.**
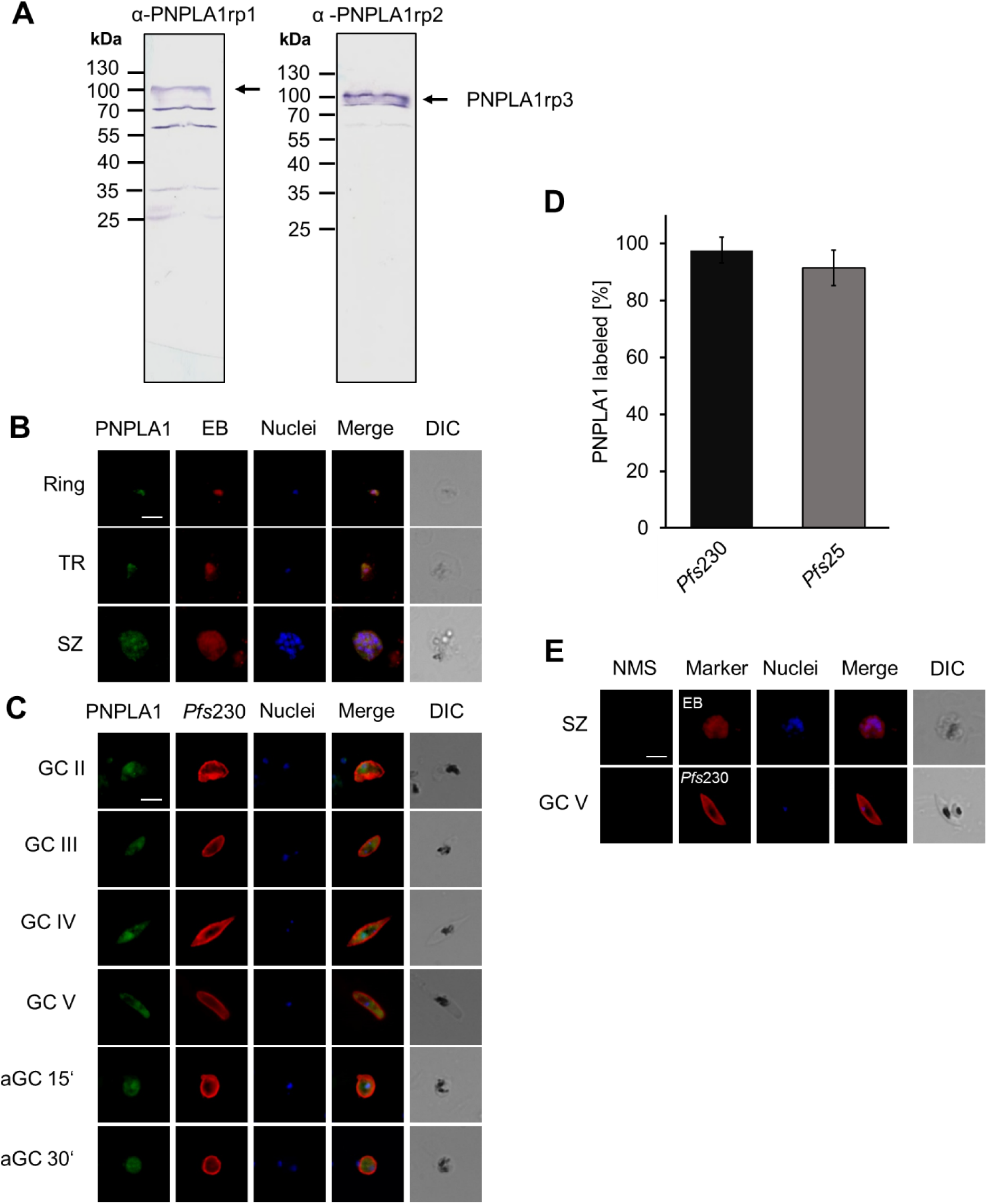
Cytosolic expression of PNPLA1 in the *P. falciparum* blood stages. **(A)** Validation of mouse antisera directed against PNPLA1rp1 and PNPLA1rp2. The PNPLA1-specific mouse antisera were employed in WB to detect the bacterially expressed full length GST-tagged PNPLA1rp3 (∼105 kDa). **(B, C)** Immunolocalization of PNPLA1 in the asexual (**B**) and (**C**) sexual blood stages of *Plasmodium falciparum.* Rings, trophozoites (TR), schizonts (SZ), gametocytes (GCII-GCV) and activated gametocytes (aGC; at 15 and 30 min p.a.) of WT were labelled with mouse anti-PNPLA1rp2 antisera (green). The asexual blood stages were stained with EB, gametocytes and activated gametocytes were counterlabelled with rabbit anti-*Pfs*230 sera (red). The parasite nuclei were highlighted by Hoechst33342 nuclear stain (blue). **(D)** Gender-specificity of PNPLA1. WT gametocytes were immunolabelled with mouse anti-PNPLA1rp1 and counterlabelled with either a rabbit antisera against *Pfs*25 (female) or *Pfs*230 (male and female). For each setting, 100 gametocytes were evaluated for co-labelling of PNPLA1. The experiment was performed in triplicate (mean ± SD). **(E)** IFA control. Schizonts (SZ) and mature gametocytes (GCV) were subjected to IFA as described above and labelled with NMS. DIC, differential interference contrast. Bar, 5 µm. Results (B-D) are representative for three independent experiments; (A) was performed once.

**FIGURE S4.**
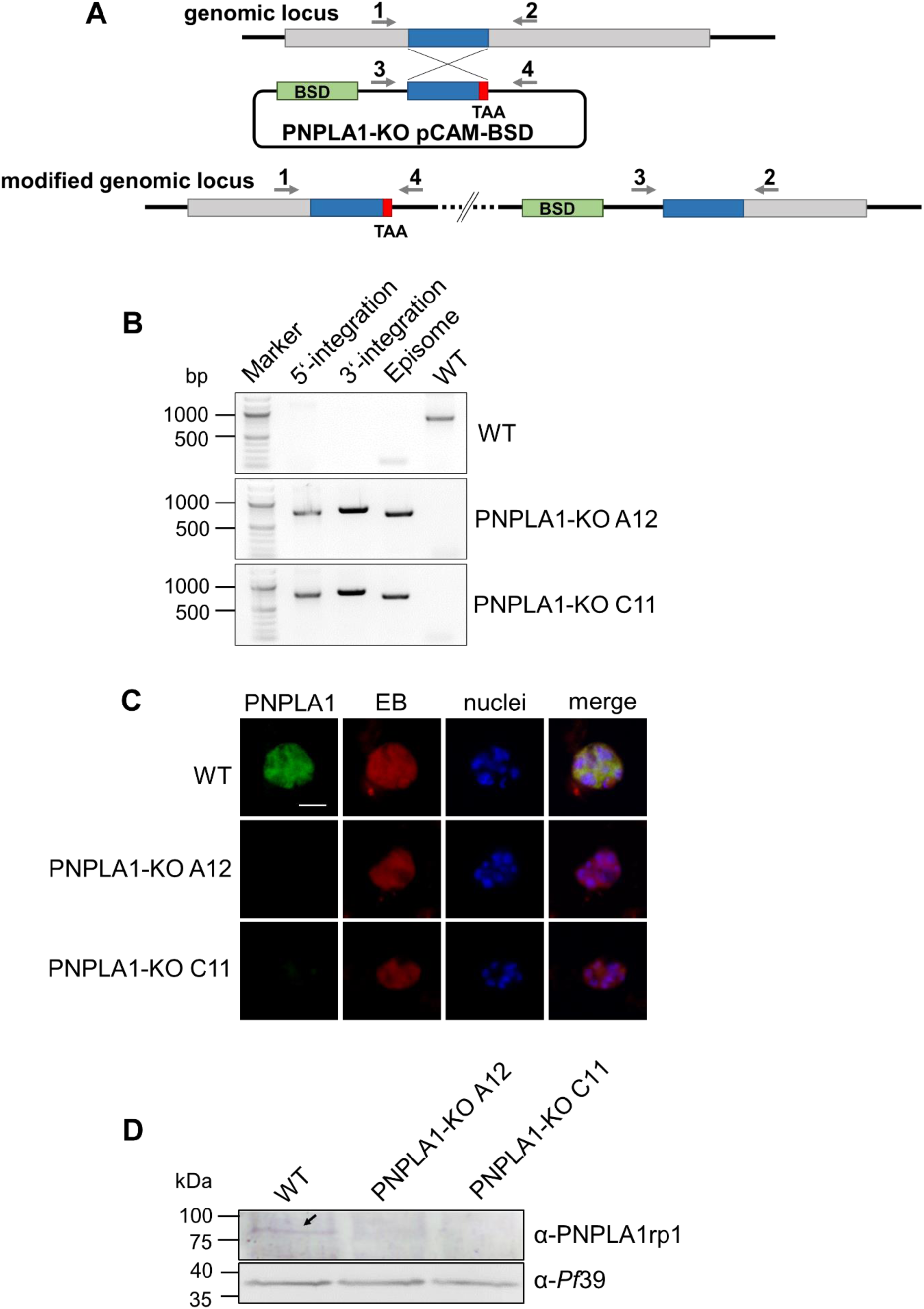
Generation of the PNPLA1-KO lines. **(A)** Schematic depicting the single-crossover homologous integration strategy to generate the *pnpla1* gene-disruptant mutants. Arrows with numbers mark the position of primers used to investigate the integration of disruption vector *pnpla1*-KO pCAM-BSD. BSD, *blasticidin-S deaminase* gene conferring resistance to blasticidin. **(B)** Confirmation of gene locus integration of the *pnpla1* pCAM-BSD vector. Diagnostic PCR on isolated gDNA of lines PNPLA1-KO A12 and C11 confirm gene locus integration of the *pnpla1*-KO pCAM-BSD vector. Primers 1 and 4 were used for amplification of the 5’-integration (773 bp) and primers 2 and 3 for 3’-integration (833 bp) of the vector. WT *pnpla1* locus was amplified with combination of primers 1 and 2 (881 bp) and episomal DNA was amplified with combination of primers 3 and 4 (712 bp). Isolated gDNA from mock control (WT) was used for negative control. **(C)** PNPLA1 detection in the PNPLA1-KO. Schizonts of WT and the PNPLA1-KO lines A12 and C11 were immunolabelled with mouse anti-PNPLA1rp1 antisera (green) and counterlabelled with EB (red). Nuclei were highlighted with Hoechst33342 nuclear stain (blue). Bar, 5 µm. **(D)** Analysis of PNPLA1 expression in the PNPLA1-KO. Lysates of schizonts of WT and the two PNPLA1-KO lines A12 and C11 were immunoblotted with mouse antisera against PNPLA1 (∼78 kDa). Equal loading was confirmed using mouse antisera against *Pf*39 (∼39 kDa). Results (B-D) are representative for three independent experiments.

**FIGURE S5.**
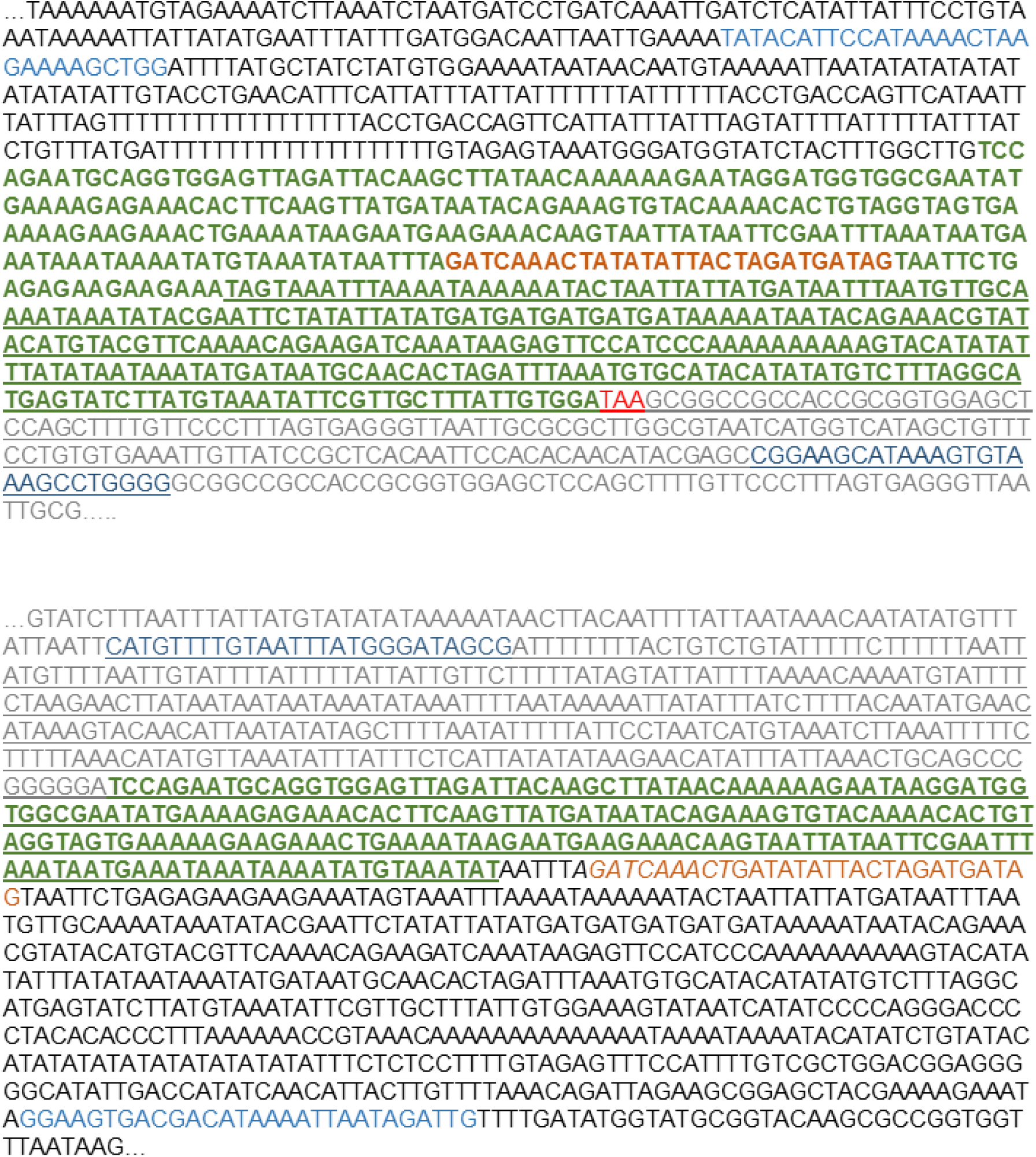
Sequencing of the integration loci of the PNPLA1-KO line A12. Sequencing of the 5’-integration locus (top) and the 3’-integration locus (bottom). The sequences corresponding to the homologous sequence on the integration vector are indicated with green letters; sequences corresponding to the WT PNPLA1 locus are indicated in black letters; sequences corresponding to the vector backbone of the pCAM-BSD vector are indicated with grey letters; sequences in blue represent primer sequences used for amplification of the sequencing locus; sequences in orange indicated sequencing primers. The stop codon is highlighted in red; underlined sequences were confirmed by sequencing. Sequences are shown in 5’-3’-orientation.

**FIGURE S6.**
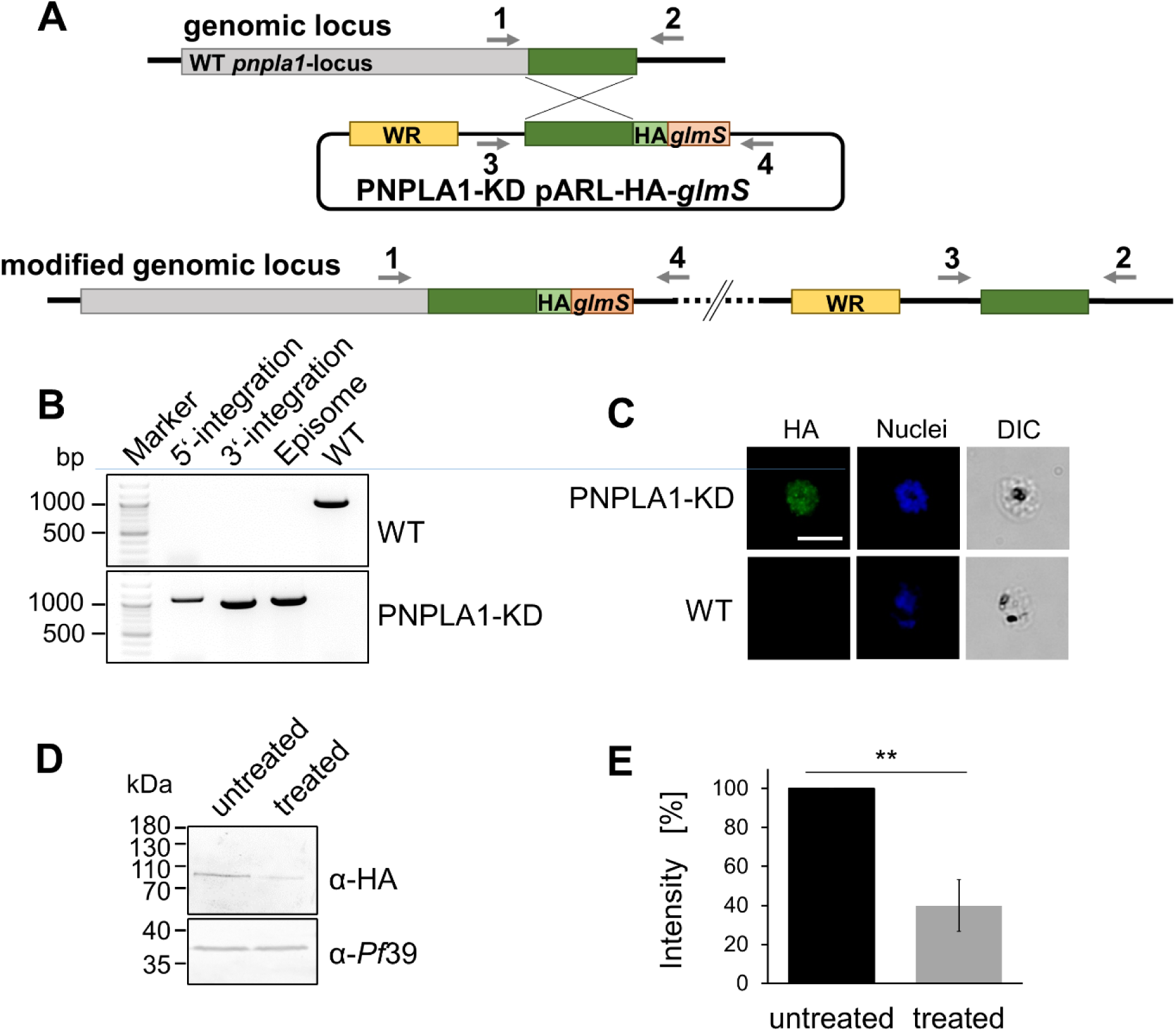
Generation of the PNPLA1-KD line. **(A)** Schematic depicting the single-crossover homologous recombination strategy for the generation of the PNPLA1-KD line. The coding region of *pnpla1* was fused at the 3’-region to a HA-encoding sequence followed by the *glmS*-ribozyme sequence. The numbered arrows indicate positions of primers used to confirm integration of the *pnpla1* pARL-HA-*glmS* vector. WR, *hDHFR* gene conferring resistance to WR99210. **(B)** Confirmation of gene locus integration of the *pnpla1* pARL-HA-*glmS* vector. Diagnostic PCR confirmed vector integration in the PNPLA1-KD parasite line. Primers 1 and 4 were used to demonstrate 5’-integration (1124 bp) and primers 2 and 3 were used to demonstrate 3’-integration (1019 bp). Primers 1 and 2 were used for amplification of the WT *pnpla1*-locus (1062 bp), while primers 3 and 4 detect the episomal plasmid (1079 bp). Isolated gDNA from mock control (WT) was used for negative control. **(C)** PNPLA1 detection in the PNPLA1-KD. Schizonts of WT and the PNPLA1-KD were immunolabeld with rabbit anti-HA antibodies (green) and nuclei were highlighted with Hoechst33342 nuclear stain (blue). DIC, differential interference contrast. Bar, 5 µm. **(D)** Analysis of PNPLA1 expression in the PNPLA1-KD. Lysates of asexual blood stages either treated with 2.5 mM GlcN for 36 h or untreated parasites were subjected to WB using rabbit anti-HA antibody to detect PNPLA1-HA (∼82 kDa). Equal loading was confirmed using mouse antisera against *Pf*39 (∼39 kDa). (**E**) Quantification of PNPLA1 expression in the PNPLA1-KD. Protein band intensities of three independent WB were quantified using ImageJ 1.51f and normalized to the respective *Pf*39 protein band intensity (mean ± SD; untreated set to 100%). **p≤0.01 (Student’s t-test). Results (B-D) are representative for three independent experiments.

**FIGURE S7.**
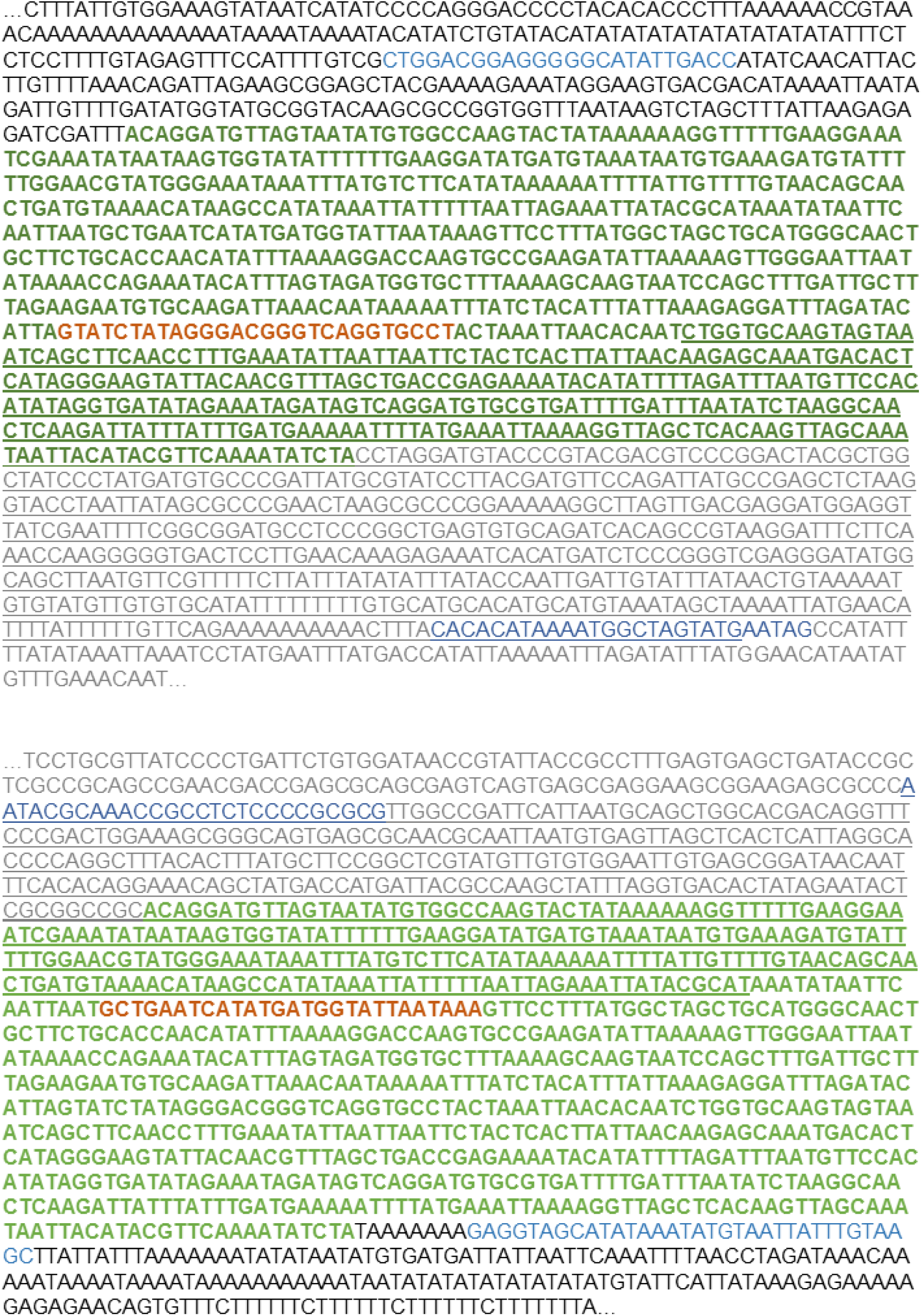
Sequencing of the integration locus of the PNPLA1-KD parasite line. Sequencing of the 5’-integration locus (top) and the 3’-integration locus (bottom). The sequences corresponding to the homologous sequence on the integration vector are indicated with green letters; sequences corresponding to the WT PNPLA1 locus are indicated in black letters; sequences corresponding to the vector backbone of the pARL-HA-*glmS* vector are indicated with grey letters; sequences in blue represent primer sequences used for amplification of the sequencing locus; sequences in orange indicate the sequencing primers; underlined sequences were confirmed by sequencing. Sequences are shown in 5’-3’-orientation.

**FIGURE S8.**
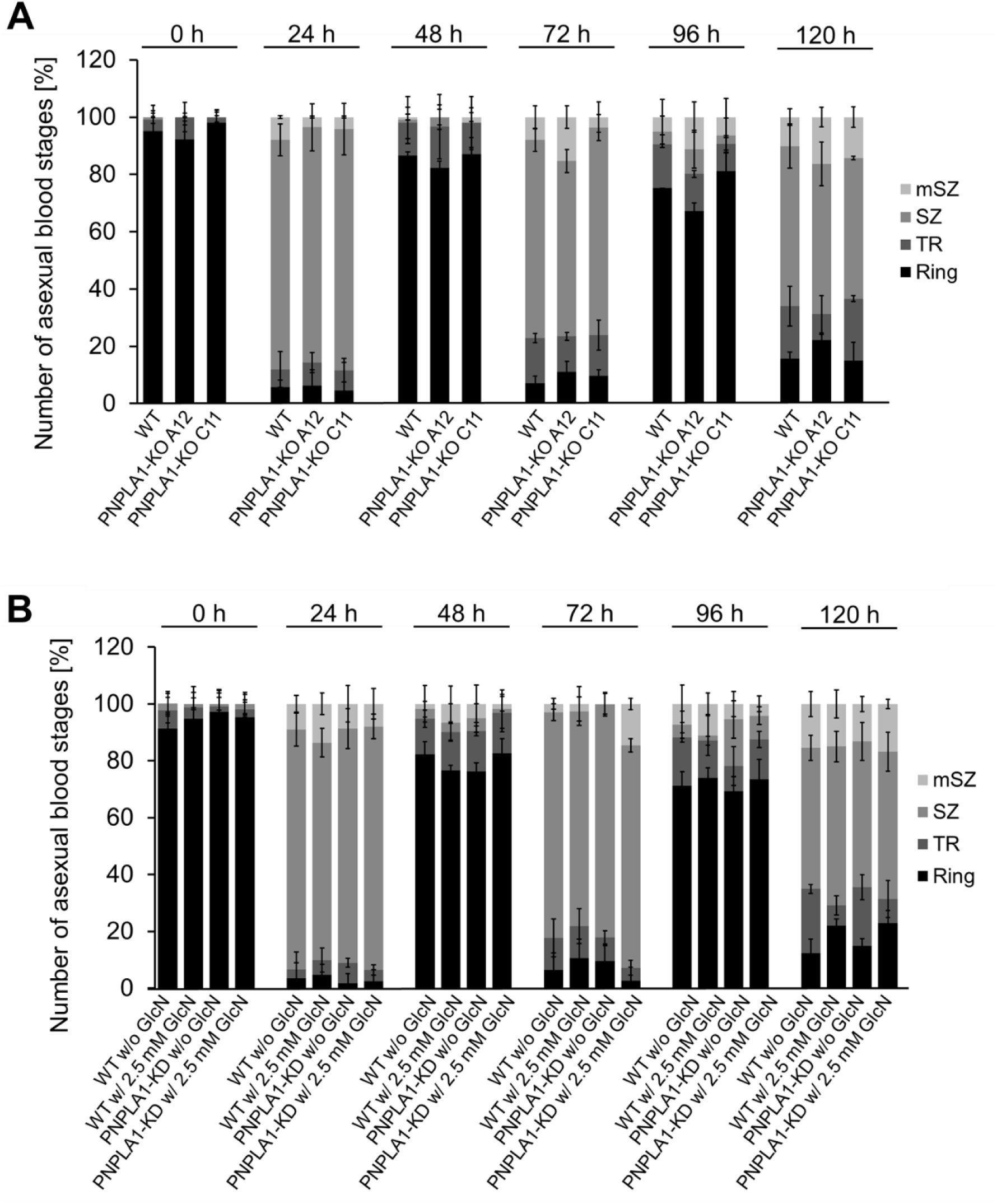
Asexual blood stage replication of the PNPLA1-deficient lines. **(A)** Synchronized ring stage cultures of WT and of PNPLA1-KO parasite lines A12 and C11 with a starting parasitemia of 0.25% were maintained in cell culture medium over a time-period of 0-120 h. Rings, trophozoites (TR), immature schizonts (SZ) and mature schizonts (mSZ), were determined in a total number of 100 infected RBCs every 24 h via Giemsa-stained blood smears. **(B)** Ring stage cultures of WT and PNPLA1-KD were maintained in cell culture medium supplemented with 2.5 mM GlcN over a time-period of 0-120 h. Untreated cultures without GlcN were used as negative control. The different developmental stages were determined as described in (A). The experiments (A, B) were performed in triplicate (mean ± SD). The results (A, B) are representative of two independent experiments.

**FIGURE S9.**
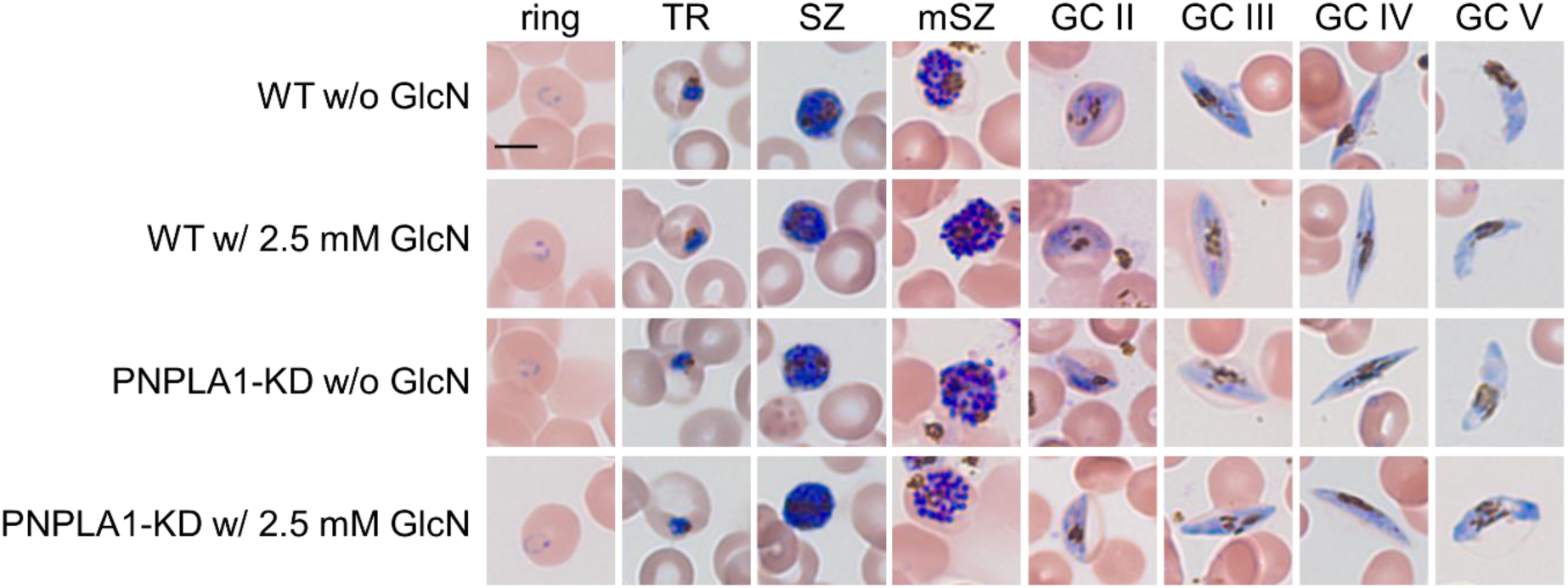
Morphology of the PNPLA1-KD blood stages. The morphology was demonstrated via Giemsa staining of asexual blood stages and gametocytes of the PNPLA1-KD and WT upon or without treatment with 2.5 GlcN. TR, trophozoites; SZ, schizonts; mSZ, mature schizonts; GC, gametocytes II-V. Bar, 5 µm.

**FIGURE S10.**
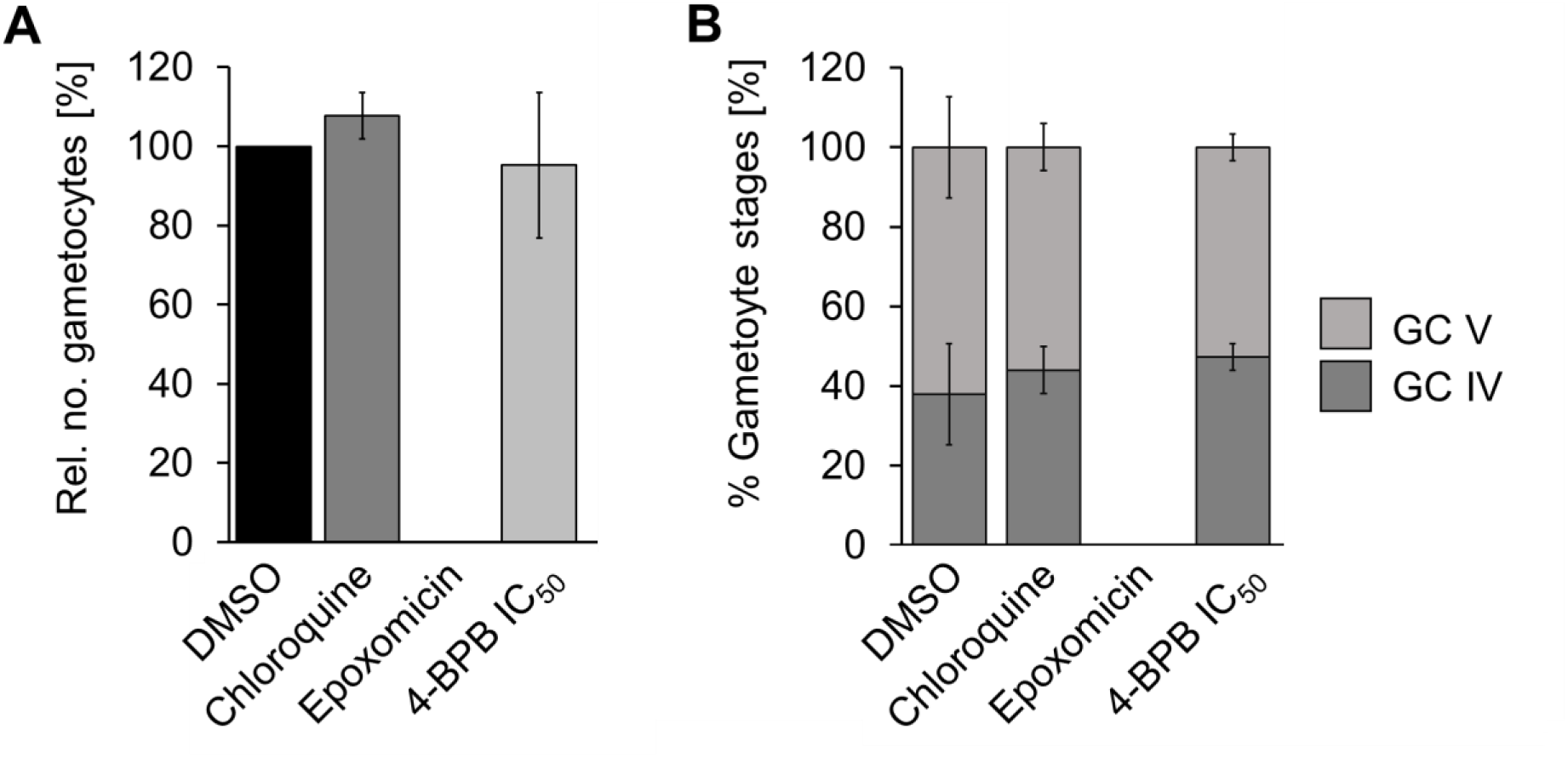
Effect of the PLA_2_ inhibitor 4-BPB on gametocyte development. The effect of 4-BPB on gametocyte development. 4-BPB at IC_50_ concentration was added to stage II gametocytes for 2 d. The numbers of total gametocytes (**A**) and the percentage of stage IV and V gametocytes (**B**) were determined at day 10 using Giemsa-stained blood smears. Epoxomicin at a concentration of 60 nM was used as positive control; 1.0% v/v DMSO and chloroquine at a concentration of 12.1 nM were used as negative controls. Experiments (A, B) were performed in triplicate (mean ± SD; DMSO set to 100%). Results (A, B) are representative of three independent experiments.

**FIGURE S11.**
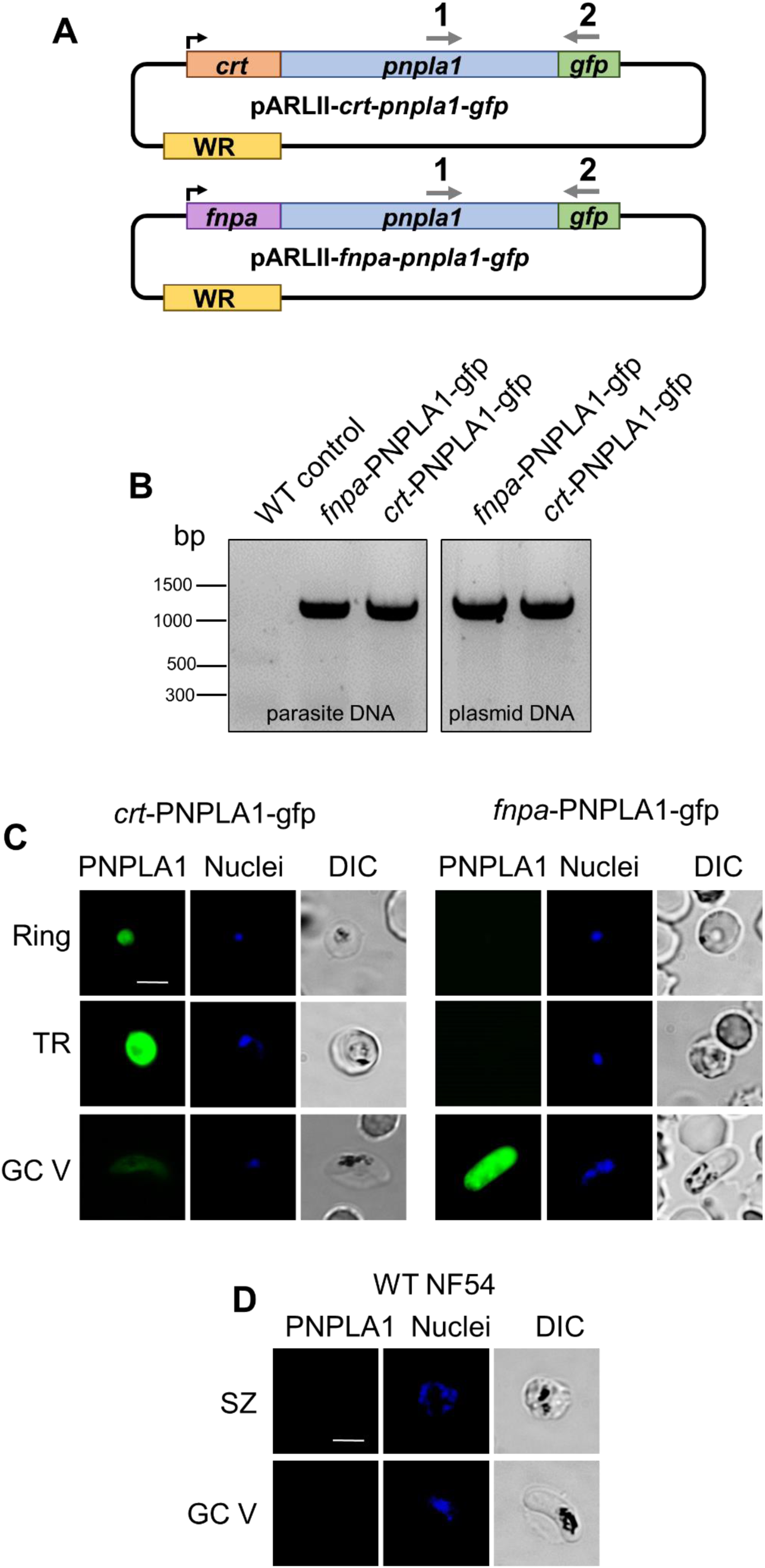
Episomal overexpression of PNPLA1-GFP. **(A)** Schematic depicting the episomal expression vectors pARLII-*crt*-*pnpla1*-*gfp* and pARLII-*fnpa-pnpla1*-*gfp*, exhibiting either the *crt* promoter or the *fnpa* promotor. **(B)** Confirmation of episomal presence in the blood stages. Diagnostic PCR on isolated gDNA from transfected lines *crt*-PNPLA1-GFP and *fnpa*-PNPLA1-GFP confirmed presence of vector DNA. Primers 1 and 2 were used to specifically amplify vector DNA (1108 bp). Purified plasmid DNA used for transfections were used as controls. **(C)** Confirmation of PNPLA1-GFP expression in the transfectants via live imaging. Transfected rings, trophozoites (TR) and mature gametocytes (GCV) of lines *crt*-PNPLA1-GFP and *fnpa*-PNPLA1-GFP were immunolabelled with mouse anti-PNPLA1rp1 antisera and nuclei were highlighted with Hoechst33342 nuclear stain. Bar, 5 µm. **(D)** IFA control. Schizonts (SZ) and mature gametocytes (GCV) were subjected to IFA as described above and labelled with sera from non-immunized mice (NMS). Bar, 5 µm. Results (B-D) are representative of three independent experiments.

**FIGURE S12.**
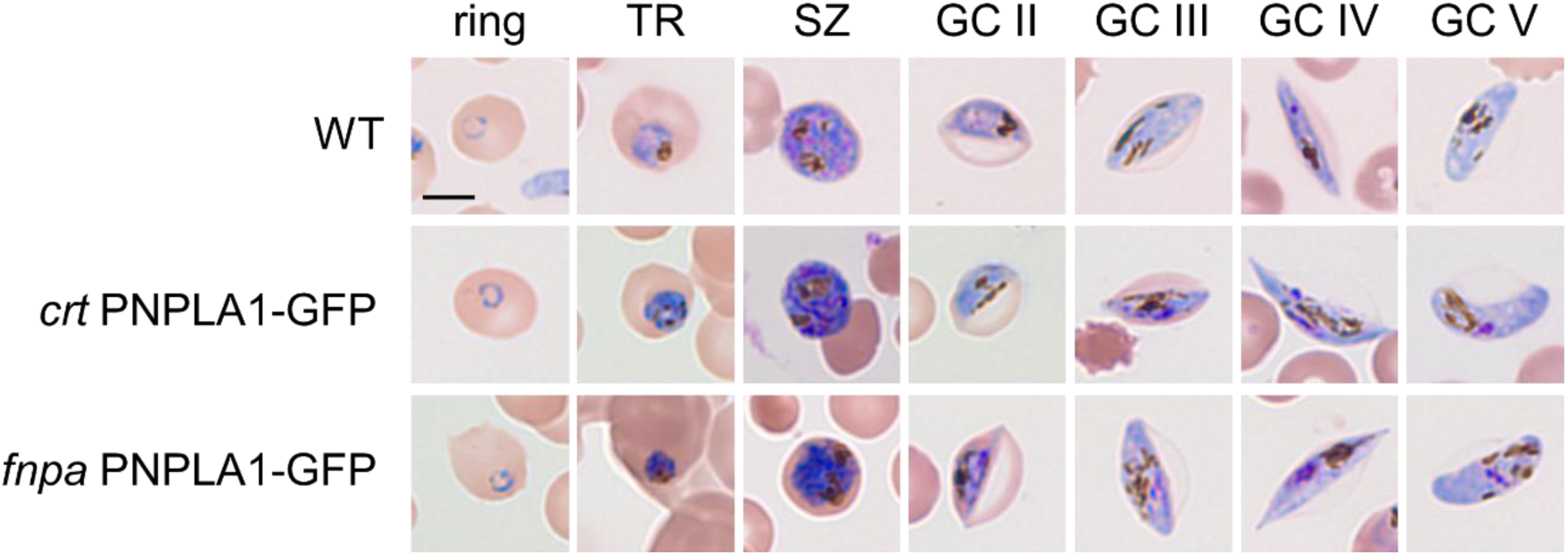
Morphology of the blood stages upon episomal overexpression of PNPLA1-GFP. The morphology was demonstrated via Giemsa staining of asexual blood stages and gametocytes of lines *crt*-PNPLA1-GFP and *fnpa*-PNPLA1-GFP and WT. TR, trophozoites; SZ, schizonts; GC, gametocytes II-V. Bar, 5 µm.

**FIGURE S13.**
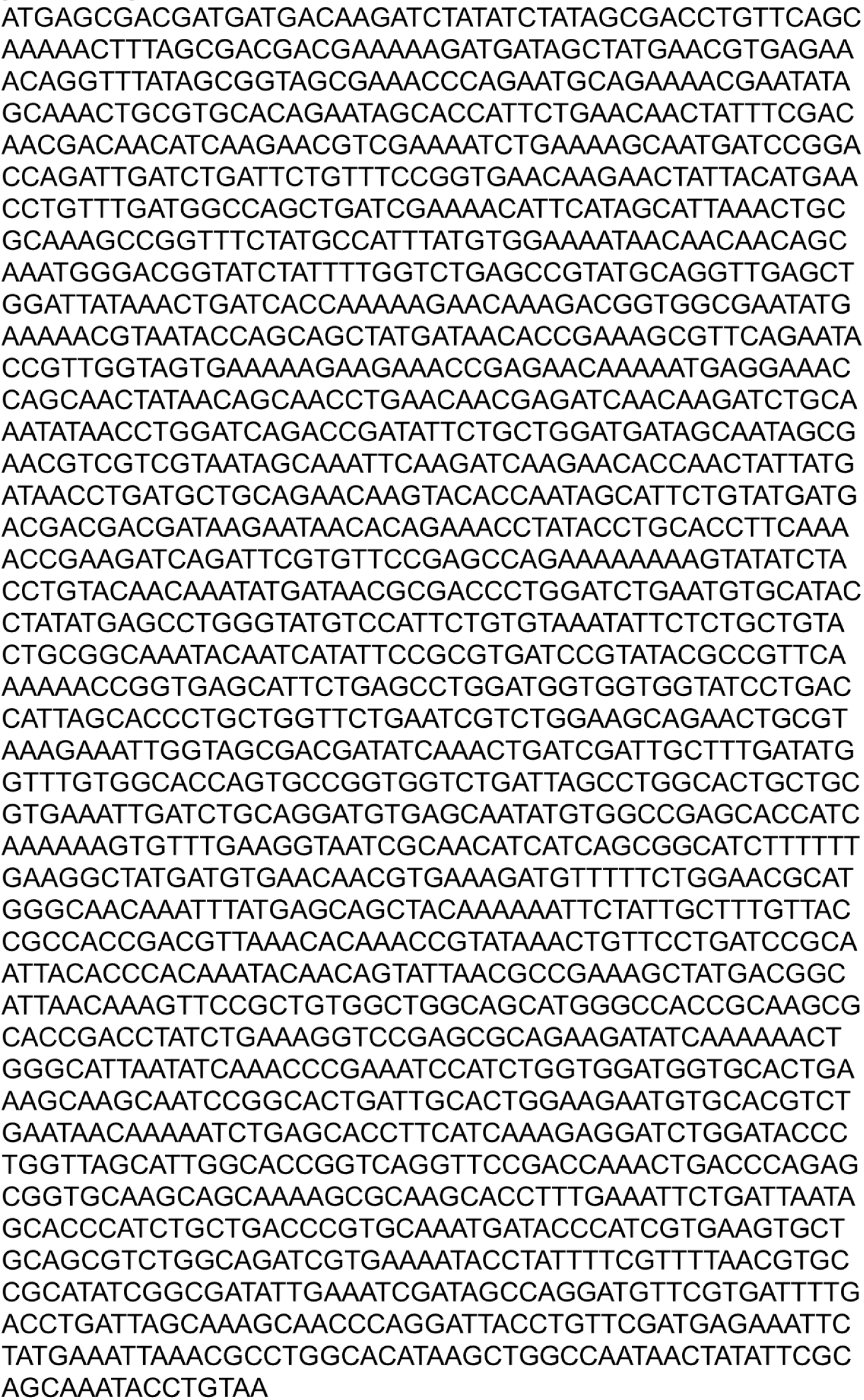
Gene sequence of PNPLA1 codon-optimized for recombinant protein expression in *E. coli*. Sequence is shown in 5’-3’-orientation.

